# G-quadruplex forming regions in *GCK* and *TM6SF2* are targets for differential DNA methylation in metabolic disease and hepatocellular carcinoma patients

**DOI:** 10.1101/2024.03.21.585382

**Authors:** Angelika Lahnsteiner, Victoria Ellmer, Anna Oberlercher, Zita Liutkeviciute, Esther Schönauer, Bernhard Paulweber, Elmar Aigner, Angela Risch

## Abstract

The alarming increase in global rates of metabolic diseases (MetDs) and their association with cancer risk renders them a considerable burden on our society. The interplay of environmental and genetic factors in causing MetDs may be reflected in DNA methylation patterns, particularly at non-canonical (non-B) DNA structures, such as G-quadruplexes (G4s) or R-loops. To gain insight into the mechanisms of MetD progression, we focused on DNA methylation and functional analyses on intragenic regions of two MetD risk genes, the glucokinase (*GCK*) exon 7 and the transmembrane 6 superfamily 2 (*TM6SF2*) intron 2-exon 3 boundary, which harbor non-B DNA motifs for G4s and R-loops.

Pyrosequencing of 148 blood samples from a nested cohort study revealed significant differential methylation in *GCK* and *TM6SF2* in MetD patients versus healthy controls. Furthermore, these regions harbor hypervariable and differentially methylated CpGs also in hepatocellular carcinoma versus normal tissue samples from The Cancer Genome Atlas (TCGA). Permanganate/S1 nuclease footprinting with direct adapter ligation (PDAL-Seq), native polyacrylamide DNA gel electrophoresis and circular dichroism (CD) spectroscopy revealed the formation of G4 structures in these regions and demonstrated that their topology and stability is affected by DNA methylation. Detailed analyses including histone marks, chromatin conformation capture data, and luciferase reporter assays, highlighted the cell-type specific regulatory function of the target regions. Based on our analyses, we hypothesize that changes in DNA methylation lead to topological changes, especially in *GCK* exon 7, and cause the activation of alternative regulatory elements or potentially play a role in alternative splicing.

Our analyses provide a new view on the mechanisms underlying the progression of MetDs and their link to hepatocellular carcinomas, unveiling non-B DNA structures as important key players already in early disease stages.

## 1. Introduction

Overweight and obesity represent a major health issue of the 21^st^ century. Their worldwide rate tripled from 1975 to 2016^1^, affecting approximately 39% of the global population^2^. Alarmingly, childhood obesity also increased from 0.7-0.9% in 1975 to 5.6-7.8% in 2016^3^. Caloric imbalance causing obesity is related to dietary habits, physical inactivity, stress, sleep patterns, social and educational status, and endocrine disorders^4^. Excessive fat accumulation increases the risk of developing metabolic diseases (MetDs) such as the metabolic syndrome (MetS), type 2 diabetes (T2D), metabolic dysfunction-associated fatty liver disease (MAFLD)^5,6^, and cardiovascular disease (CVD)^7^. Furthermore, MetD patients have an elevated risk for gastric^8^, pancreatic^9^, hepatocellular^10^, colorectal^11^, breast^12^ and lung cancer^13,14^.

MAFLD is the most common form of chronic liver disease in industrialized societies^15^ with a worldwide prevalence of 25%^16^, and is strongly associated with T2D^17^. One hallmark of MAFLD is the development of large cytoplasmic lipid droplets in hepatocytes with the accumulation of proteins involved in lipid homeostasis on their surface. The key disease mechanism in MAFLD is very likely related to peripheral insulin resistance^17,18^. In addition, fatty acid β-oxidation is impaired in MAFLD owing to mitochondrial respiratory chain dysfunction, which, in combination with the accumulation of incompletely metabolized fatty acids, contributes to oxidative and lipotoxic stress in hepatocytes. This facilitates the progression from simple steatosis to advanced stages of MAFLD which can further progress to liver fibrosis, cirrhosis, and to hepatocellular carcinoma (HCC)^19^. Currently, we do not fully understand the molecular mechanisms leading to the development and progression of MAFLD, but they clearly involve a complex interplay between genetic predisposition and environmental factors influencing epigenetic processes, such as DNA methylation at cytosines (5-methylcytosine, 5mC). Studies investigating 5mC have identified differentially methylated positions (DMPs) associated with hepatic fat or key signaling pathways regulating glycolysis and *de novo* lipogenesis^20–24^. However, a mechanistic explanation for the impact of those detected DMPs on cellular processes is often lacking. Particularly, if these DMPs are located in intergenic regions or in gene bodies without overlapping regulatory elements, interpreting their biological functions remains difficult.

New insights arise from the observation that G-rich sequences, with the ability to form non-canonical (non-B) DNA structures called G-quadruplexes (G4s), are sites of methylation instability^25^, and actively influence DNA methylation^26^. G4s are formed via the consensus motif G_≥3_N_x_G_≥3_N_x_G_≥3_N_x_G_≥3_ and can adopt intra- or intermolecular planar G4 tetrads held together by Hoogsteen hydrogen bonds with a 1-7 bp long linker of any base (N) and stabilized by a monovalent cation, *in vivo* typically potassium^27–29^. These structures are found in regulatory regions such as promoters^30,31^, enhancers^32^, or near splice sites^33^, whereas they are less often detected in exons and on the template strand^34^. They often co-occur with RNA:DNA hybrids (R-loops), formed during the transcription of long non-coding (lncRNAs) on the displaced strand and each stabilizing the other^35^. G4s act as binding hubs for several transcription factors (TFs)^36,37^ and DNA methyltransferases (DNMTs)^26,38,39^, which is underlined by their manifold biological roles extensively reviewed elsewhere^40–42^.

G4s are fairly sensitive to changes in ion concentrations like potassium^43^, the pH^44^, or glucose concentrations^45^. Moreover, DNA methylation influences the thermodynamic properties of the DNA duplex ^46^ and therefore, methylation of CpGs within a G4-forming sequence (G4-CpGs) possibly triggers its rearrangement from a DNA duplex to a non-B DNA structure. This is supported by the observation of enhanced G4 formation due to DNA methylation within the first exon of the human reverse transcriptase (hTERT) gene, which in turn, leads to repressed gene transcription ^47^. Furthermore, DNA methylation leads to increased stability of G4 structures in the *BCL2*^48^, *VEGF*^49^ and *FMR1*^50^, whereas decreased stability was detected for the *MEST* G4^51^ in *in vitro* assays. Even changes in the G4 topology from parallel to antiparallel in *c-KIT,* or a complete loss of G4 formation in *HRAS* can be caused by DNA methylation^52^. Genome-wide approaches revealed that methylation of CpGs within G4s (G4-CpGs) is significantly decreased compared to CpGs outside of G4s, especially among those CpGs located in CpG islands (CGIs)^26,53^. Furthermore, it has been shown, that DNMT1 preferentially binds to G4 structures rather than to hemi-methylated B-DNA, which results in the inhibition of the DNMT1 and local hypomethylation at the G4 structure^26^. There are still contradictory discussions, since the mentioned studies mostly have shown that G4-CpGs are targets of hypomethylation, while a recent work investigating DNA methylation of aging clocks suggests a general DNA methylation instability at G4-CpGs, rather than a particular hypo- or hypermethylation of these G4-CpGs^25^. Furthermore, whereas studies investigating individual G4 motifs have shown that DNA methylation can also stabilize G4s, genome-wide analysis demonstrated that the stability of G4s with a maximum distance of 50 nucleotides to the CGI is associated with hypomethylation and in cases where the G4 containing CGI is located in closed and hypermethylated chromatin, the G4 stability was lowest^54^. Most analyses focused on promoters and on regions overlapping CGIs, but a considerable number of regulatory regions, like alternative promoters^55,56^ or enhancers^57^, occur outside of CGIs where different regulatory mechanisms probably apply. Selective DNA methylation loss in CG-poor regions can make chromatin accessible for alternative transcription, particularly in HCC^58^, and we have previously associated differential methylation with alterations in promoter activation^59^. This indicates that we are still in the early stages of understanding the mechanisms involved, and that different genetic locations probably underlie distinct mechanisms. Therefore, alterations in the G4 landscape could alter transcription factor binding and activate alternative regulatory elements or alternative splicing events, causing aberrant gene expression patterns that contribute to MAFLD and may even promote the development of more severe diseases such as hepatocellular carcinoma.

Patients suffering from metabolic diseases are at a higher risk of developing cancer during their lifetime. Therefore, it is necessary to understand which processes are involved and DNA methylation at G4s could be one of these mechanisms, probably already arising in a noncancerous but dysregulated metabolic stage. Hence, we aimed to analyze DNA methylation and G4 formation in MetD patients and in HCC in two intragenic regions outside of CGIs with annotated regulatory elements, as well as the occurrence of putative G4 forming motifs and lncRNAs in the key metabolism genes glucokinase (*GCK*) and transmembrane 6 superfamily 2 (*TM6SF2*). *GCK* deficiency has been linked with T2D^60^ and neonatal diabetes^61^, while a common *GCK* SNP has been associated with hyperglycemia^62^ and several alternative *GCK* promoters are annotated for the gene body and differential methylation of the *GCK* CGI spanning exons 9-10 had been linked with CVD^63^. The intron 2-exon 3 boundary in *TM6SF2* overlaps with a prognostic lncRNA for cancer^64,65^, it shows many different transcripts and rs58542926 C>T, the SNP prognostic for MAFLD^66^ and HCC^67^, is located downstream of the annotated regulatory region. rs58542926 C>T is routinely checked in the clinics and potentially could have an impact on the methylation of the next regulatory element.

Herein, we report differential DNA methylation in exon 7 of *GCK* and within the intron 2-exon 3 boundary of *TM6SF2* in blood samples from a MetD patient cohort in G4 forming regions, which are involved in the activation of alternative regulatory elements and possibly also in alternative splicing. These new insights emphasize the critical need to understand the interplay between aberrant DNA methylation and non-B DNA structure formation occurring already at an early disease stage.

## 2. Results

### Cohort demographics and description

To investigate DNA methylation in putative intragenic G4-forming regions in *GCK* and *TM6SF2*, we performed locus-specific 5mC analysis on the Qiagen Pyromark Q24 Platform for 148 blood donors which were classified as lean-healthy (LH), obese-healthy (OH), lean-MAFLD (LM), obese-MAFLD (OM), and advanced MetD (aMetD, including MAFLD and poorly controlled T2DM patients) according to their metabolic parameters, as described in the *Methods* section. The mean values of the clinical parameters are provided in Table 1 and Supplement Fig. 1. The age of our study cohort ranged from 42-77 years, albeit the OM and aMetD groups showed significantly elevated age compared to the LH group (LH vs. OM *p*=.0152, LH vs aMetD *p*= 1.53e-05, Student’s t-test, and Supplement Table 1). This trend was not unexpected since MAFLD and T2D are known as aging-related diseases occurring at a higher frequency in the elderly (reviewed in^68^). Fasting blood glucose, insulin, and aspartate transferase (AST) levels were significantly increased in OH, OM and aMetD, but not in LM compared to LH (Students t-test, Supplement Table 1). No significant change was detected in total cholesterol levels in any group compared to healthy controls. High-density lipoprotein (HDL) was significantly decreased in OM and aMetD, and low-density lipoprotein (LDL) was only decreased in aMetD, whereas total triglycerides were increased in LM, OM, and aMetD. The detailed *p*-values are shown in the Supplement Table 1.

**Table 1.**
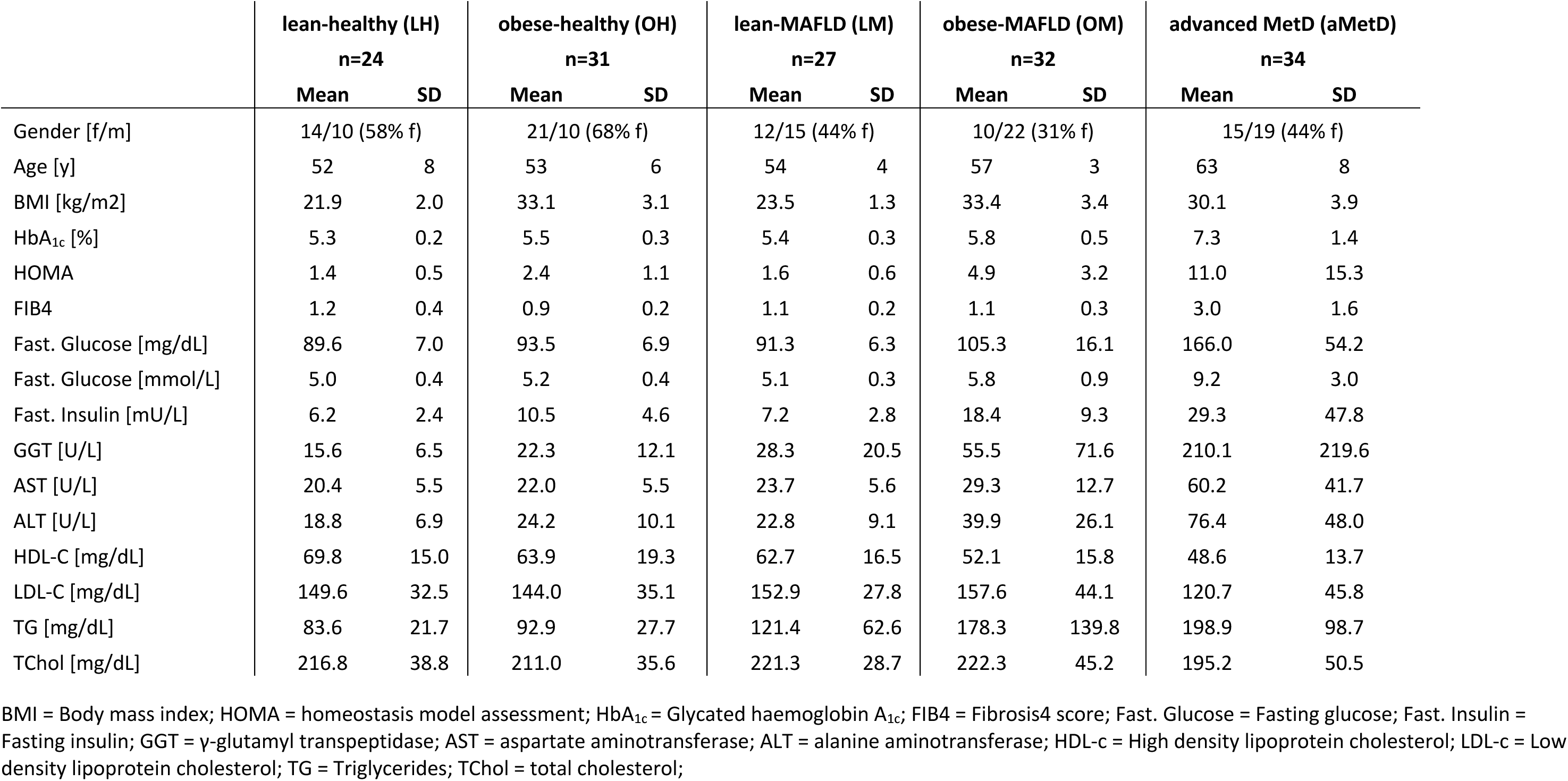
Study characteristics. Data are expressed as means ± standard deviation (SD)

|  | lean-healthy (LH)<br>n=24 |  | obese-healthy (OH)<br>n=31 |  | lean-MAFLD (LM)<br>n=27 |  | obese-MAFLD (OM)<br>n=32 |  | advanced MetD (aMetD)<br>n=34 |  |
| --- | --- | --- | --- | --- | --- | --- | --- | --- | --- | --- |
|  | Mean | SD | Mean | SD | Mean | SD | Mean | SD | Mean | SD |
| Gender [f/m] | 14/10 (58% f) |  | 21/10 (68% f) |  | 12/15 (44% f) |  | 10/22 (31% f) |  | 15/19 (44% f) |  |
| Age [y] | 52 | 8 | 53 | 6 | 54 | 4 | 57 | 3 | 63 | 8 |
| BMI [kg/m<sup>2</sup>] | 21.9 | 2.0 | 33.1 | 3.1 | 23.5 | 1.3 | 33.4 | 3.4 | 30.1 | 3.9 |
| HbA<sub>1c</sub> [%] | 5.3 | 0.2 | 5.5 | 0.3 | 5.4 | 0.3 | 5.8 | 0.5 | 7.3 | 1.4 |
| HOMA | 1.4 | 0.5 | 2.4 | 1.1 | 1.6 | 0.6 | 4.9 | 3.2 | 11.0 | 15.3 |
| FIB4 | 1.2 | 0.4 | 0.9 | 0.2 | 1.1 | 0.2 | 1.1 | 0.3 | 3.0 | 1.6 |
| Fast. Glucose [mg/dL] | 89.6 | 7.0 | 93.5 | 6.9 | 91.3 | 6.3 | 105.3 | 16.1 | 166.0 | 54.2 |
| Fast. Glucose [mmol/L] | 5.0 | 0.4 | 5.2 | 0.4 | 5.1 | 0.3 | 5.8 | 0.9 | 9.2 | 3.0 |
| Fast. Insulin [mU/L] | 6.2 | 2.4 | 10.5 | 4.6 | 7.2 | 2.8 | 18.4 | 9.3 | 29.3 | 47.8 |
| GGT [U/L] | 15.6 | 6.5 | 22.3 | 12.1 | 28.3 | 20.5 | 55.5 | 71.6 | 210.1 | 219.6 |
| AST [U/L] | 20.4 | 5.5 | 22.0 | 5.5 | 23.7 | 5.6 | 29.3 | 12.7 | 60.2 | 41.7 |
| ALT [U/L] | 18.8 | 6.9 | 24.2 | 10.1 | 22.8 | 9.1 | 39.9 | 26.1 | 76.4 | 48.0 |
| HDL-C [mg/dL] | 69.8 | 15.0 | 63.9 | 19.3 | 62.7 | 16.5 | 52.1 | 15.8 | 48.6 | 13.7 |
| LDL-C [mg/dL] | 149.6 | 32.5 | 144.0 | 35.1 | 152.9 | 27.8 | 157.6 | 44.1 | 120.7 | 45.8 |
| TG [mg/dL] | 83.6 | 21.7 | 92.9 | 27.7 | 121.4 | 62.6 | 178.3 | 139.8 | 198.9 | 98.7 |
| TChol [mg/dL] | 216.8 | 38.8 | 211.0 | 35.6 | 221.3 | 28.7 | 222.3 | 45.2 | 195.2 | 50.5 |

BMI = Body mass index; HOMA = homeostasis model assessment; HbA<sub>1c</sub> = Glycated haemoglobin A<sub>1c</sub>; FIB4 = Fibrosis4 score; Fast. Glucose = Fasting glucose; Fast. Insulin = Fasting insulin; GGT = γ-glutamyl transpeptidase; AST = aspartate aminotransferase; ALT = alanine aminotransferase; HDL-c = High density lipoprotein cholesterol; LDL-c = Low density lipoprotein cholesterol; TG = Triglycerides; TChol = total cholesterol;

### *GCK* exon 7 and the *TM6SF2* intron 2-exon 3 boundary are differentially methylated in MetD patients and are hypervariable in HCC

*GCK* is encoded on the minus strand of chr. 7 and consists of ten exons (Fig. 1A). 14 different transcripts are annotated for *GCK* in the ENSEMBL genome browser^69^. The investigated region is located in exon 7 (chr7:44,187,339-44,187,380; hg19) and is overlapped by an antisense lncRNA LOC105375258 (XR_927223.3) on the plus strand. Genome segmentation by automating chromatin state discovery and characterization (ChromHMM) in the UCSC genome browser predicts a “weak enhancer or open chromatin *cis* regulatory element” in hepatoblastoma cell line HepG2 and a “promoter region including TSS” in the lymphoblast cell line K562 (Fig. 1A, yellow rectangle)^70^. Furthermore, several alternative promoters are found in intron 7 and 8, as well as an EGR1 TF binding site (Fig. 1A, green rectangle)^70^. Significant hypomethylation was detected in CpG 1 and 2 in aMetD patients, and in CpG 3 in LM, OM and aMetD compared to LH subjects. No changes were detected in CpG 4 (Fig. 1A lower panel and Supplement Fig. 2A). In particular, a gradual reduction in DNA methylation was detected for CpG 3 with the individual disease stages, and the advanced MetD patients showing the lowest methylation.

**Figure 1.**
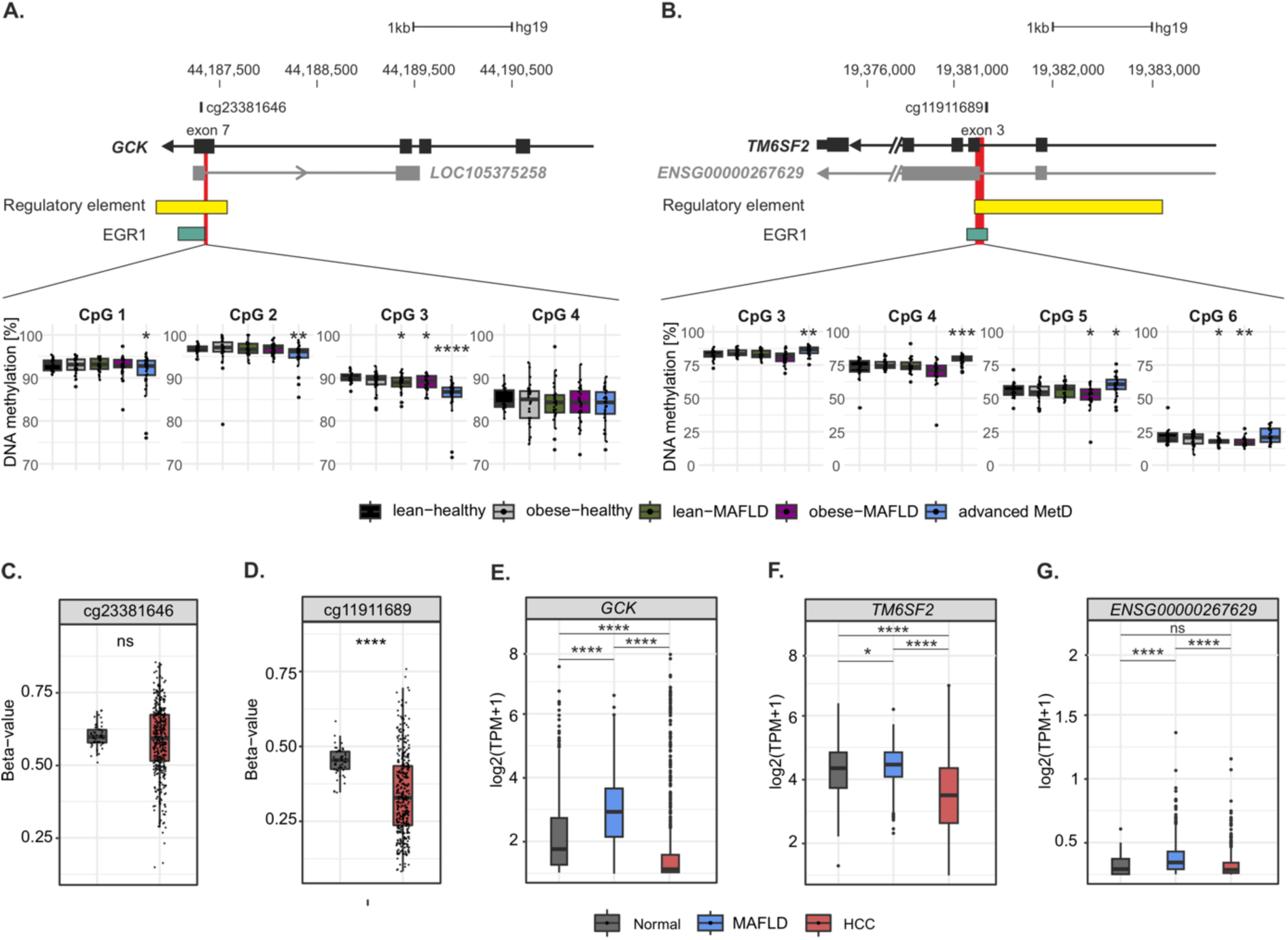
DNA methylation in *GCK* exon 7 and within the *TM6SF2* intron 2-exon 3 boundary. Detailed information about the differentially methylated regions in **A.** *GCK* and **B.** *TM6SF2* including the nearest annotated CpG positions on the Illumina Methylation array. The yellow rectangle indicates the genome segmentation by ChromHMM in UCSC genome browser^70^. The green rectangle represents the EGR1 binding motif. The pyrosequencing region is marked in red. The lower panel shows DNA methylation at the single CpG resolution by metabolic status in **A.** for *GCK* CpG 1-4 and **B.** *TM6SF2* CpG 3-6. DNA methylation data of the nearest annotated CpG on the Illumina Methylation array for **C.** *GCK* cg23381646 and **D.** *TM6SF2* cg11911689 are shown comparing HCC to normal tissue from TCGA^71^. Gene expression for **E.** *GCK*, **F.** *TM6SF2* and **G.** the *TM6SF2*-overlapping lncRNA ENSG00000267629 were obtained from the liver expression atlas (GepLiver)^72^. Pairwise comparison using Students t-test against LH (**A.-B.**) and Wilcoxon test (**C.-G.**), \**p*<.05, ** *p*<.01, *** *p*<.001, **** *p*<.0001.

In line with *GCK* exon 7, we found differential methylation also for the *TM6SF2* intron 2-exon 3 boundary (chr19:19,381,224-19,381,297; hg19, Fig. 1B). *TM6SF2* is encoded by ten exons on the minus strand of chr. 19. In total, there are five transcripts annotated, of which two result in a retained intron and one in a skipped exon overlapping our target region^69^. Notably, exon 3-5 are targets of alternative splicing events and exon 3 is annotated as cassette exon (Supplement Fig. 3)^70^. Furthermore, *TM6SF2* overlaps with the sense lncRNA ENSG00000267629 (AC138430.4), sharing the same transcription start site. This lncRNA represents a readthrough between *TM6SF2* and *HAPLN4*. ChromHMM predicts a “weak enhancer or open chromatin *cis* regulatory element” in HepG2 and a “promoter region including TSS” in K562 in the UCSC genome browser^70^, as well as an EGR1 binding site (Fig. 1B). 5mC analyses revealed that CpG 3-5 were significantly hypermethylated in the advanced MetD group. In contrast, CpG 4 and 5 were significantly hypomethylated in OM compared to LH and CpG 5 was also significantly hypomethylated in LM compared to LH (Student’s t-test, Fig. 1B lower panel and Supplement Fig. 2B).

Given that alterations in DNA methylation are also associated with aging^73^ and metabolic diseases occur at a higher frequency with increasing age, we analyzed the magnitude of age-relation compared to published aging markers located in *EDARADD* (cg09809672), *IPO8* (cg19722847), *NHLRC1* (cg22736354), *P2RXL1* (cg05442902) and *SCGN* (cg06493994). These five markers were selected based on their consistent identification in three distinct epigenetic clock studies utilizing Illumina Methylation arrays^73–75^. Consistent with published data, we detected a significant positive (*NHLRC1* and *SCGN*) and negative (*IPO8*, *EDARADD* and *P2RXL1*) association between DNA methylation and chronological age in our cohort (Supplement Fig. 4). In addition, the correlation analysis showed a weak but significant age-dependency for *GCK* CpG 3 and *TM6SF2* CpG 1, 3 and strongest for CpG 5, as well as for the mean DNA methylation (mean of CpG 1-6). Therefore, we restricted our analysis to 42–60-year-old subjects. We still observed a significant decreased DNA methylation at *GCK* CpG 3 in all groups compared to lean-healthy subjects, which reached significance even in obese-healthy patients, while this was not the case when analyzing the entire age-spectrum (Supplement Fig. 2C). *TM6SF2* CpG 3, 4 and 6 where still significantly differentially methylated, but the effect was lost in CpG 5 in 42–60-year-old subjects (Supplement Fig. 2D). This points out that aberrant DNA methylation in these regions is only partly caused by aging processes and underlies also a disease-related mechanism.

Next, we were curious if the SNP rs58542926 C>T, which causes a p.Glu167Lys (E167K) substitution in *TM6SF2*, could influence the DNA methylation in this specific regulatory region. The SNP is located 1,696 bp away from CpG 4 in exon 6. Although the risk associated variant T is more common in the advanced MetD group (Supplement Fig. 5A), we found no significant difference in DNA methylation by genotype in our study cohort (Supplement Fig. 5B). Therefore, we can exclude that the genetic variant rs58542926 C>T causes the detected differential DNA methylation within *TM6SF2* intron 2-exon 3.

Since metabolic diseases are associated with an increased risk for HCC^10^, we investigated 5mC of *GCK* and *TM6SF2* in data of *The Cancer Genome Atlas* (TCGA) from HCC compared to normal tissue^71^. The nearest annotated CpG for *GCK* covered by the Illumina Methylation array is cg23381646 located 45 bp upstream of CpG 3 (Fig. 1A). Although DNA methylation at cg23381646 was not significantly different (Wilcoxon test, Benjamini-Hochberg (BH) adjusted *p*=.58, Fig. 1C), it appeared as highly variable in tumor tissue with DNA methylation beta-values ranging from approximately 0.25-0.8.

For *TM6SF2*, TCGA data showed that the nearest annotated CpG cg11911689 (78 bp downstream of CpG 4, Fig. 1B) was significantly hypomethylated in HCC compared to normal tissue (Wilcoxon test, BH adjusted *p*=4.5E-10, Fig. 1D), and again showed a highly variable DNA methylation range in cancer compared to normal control tissue.

To explore gene expression of the target genes, we analyzed *GCK* and *TM6SF2* expression in MAFLD and HCC in the liver expression database (GepLiver)^72^, since transcriptome data were not available for the MetD cohort due to the lack of liver material. *GCK* expression was increased by 1.73-fold (Wilcoxon test, BH adjusted *p*=2.52E-32) in MAFLD, but decreased expression was detected in HCC compared to normal liver tissue (0.67-fold, Wilcoxon test, BH adjusted *p*=2.73E-32, Fig. 1E). Significant increased expression was also detected for *TM6SF2* in MAFLD (1.04-fold, Wilcoxon test, BH adjusted *p*=.035), which was again decreased in HCC by 0.76-fold (Wilcoxon test, BH adjusted *p*=7.48E-28, Fig. 1F). Of note, the expression of the *TM6SF2* overlapping lncRNA ENSG00000267629 was increased in MAFLD by 1.92-fold (Wilcoxon test, BH adjusted *p*= 4.31E-7). Although it remained slightly overexpressed in HCC, it did not reach significance (1.08-fold, Wilcoxon test, BH adjusted *p*= .58, Fig. 1G).

### Differentially methylated regions detected in *GCK* and *TM6SF2* are able to form non-B DNA structures

To investigate if the detected DMPs are located in non-B DNA structures and if they are formed in living cells, we retrieved information about R-loop forming sequences (RLFS)^76^ and putative G4 motifs (PQS) from pqsfinder^77^. The *GCK* DMR harbors several G4 motifs, but no predicted RLFS, albeit this exon is overlapped by the annotated antisense lncRNA LOC105375258 (Fig. 2A left panel). All four analyzed CpG positions in *GCK* exon 7 are located in a G4 motif in the non-template strand, with the top differentially methylated CpG 3 located in the longest loop of the G4 between the G-run two and three. Notably, the G4 includes a bulge, represented as an interruption in the G-run, between the third and fourth G of G-run two (Sequence motif in Fig. 2A left panel). Furthermore, the downstream exons 8-10 harbor both RLFS and PQS (Fig. 2A left panel). DNA methylation of this downstream region was investigated elsewhere and showed also differential methylation^63^. The *TM6SF2* DMR contained a RLFS as well as PQS (Fig. 2A middle panel). The differentially methylated CpGs 4-6 were located directly within the predicted G4 motif in the template strand and CpG 1-3 flanked the G4 motif. As negative control region we have used an intronic region of *PARD3B* without annotated PQS or RLFS (Fig. 2A right panel). *In vitro* G4 formation was verified by analyzing published G4-sequencing (G4-Seq) maps based on a polymerase stop assay in a buffer containing the G4 stabilizing ligand pyridostatin (PDS) in HEK293T cells (Fig. 2A, black track)^78^. Strong signal intensities in *GCK* and *TM6SF2,* but not in *PARD3B*, confirmed the ability of the predicted motifs to form DNA secondary structures. As G4s are not necessarily formed in living cells and their formation varies strongly in different cell types, we analyzed published G4 ChIP-Seq data in HEK293 cells^79^. Especially, *GCK* shows enriched signal intensities compared to its input control over the whole region spanning from exon 7 to exon 10, rather than distinct peaks similar to the G4-Seq signals (Fig. 2A left panel). *TM6SF2* showed a more discrete peak for replicate 1 but not for replicate 2 (Fig. 2A middle panel). Since, we could not infer a clear picture for G4 formation from the antibody-based ChIP-Seq data, particularly for *GCK* exon 7, we aimed to establish an alternative chemical method based on permanganate/S1 nuclease footprinting^80^ which in general detects regions of single-stranded DNA (ssDNA). We updated this method by linking it to direct adapter ligation on streptavidin-coated beads, followed by an on-bead PCR published as PDAL-Seq (<u>p</u>ermanganate/S1 nuclease footprinting with <u>d</u>irect <u>a</u>dapter ligation and <u>seq</u>uencing)^81^. Since PDAL-Seq is a footprinting method, we expected low or absent read frequencies directly at the non-B DNA structure flanked by increased read frequencies at the non-B DNA borders. For HEK293 cells, we did not detect a direct overlap of PDAL-footprints with G4-Seq peaks at *GCK* exon 7 rather than a slight upstream shift, but in MCF7 cells the non-B-footprint was more obvious, highlighting the cell type differences (Fig. 2A left panel, light blue highlighted box). Of note, footprints detected in *GCK* exon 8-10 were more pronounced and overlapped well with the strong G4-Seq signals, while the peaks from the antibody-based G4 ChIP-Seq method were less clear (Fig. 2A, left panel, light red highlighted box). In contrast, the intronic region of *PARD3B* did not contain a G4 motif, nor did it form G4s *in vitro* or in living cells (Fig. 2A right panel).

**Figure 2.**
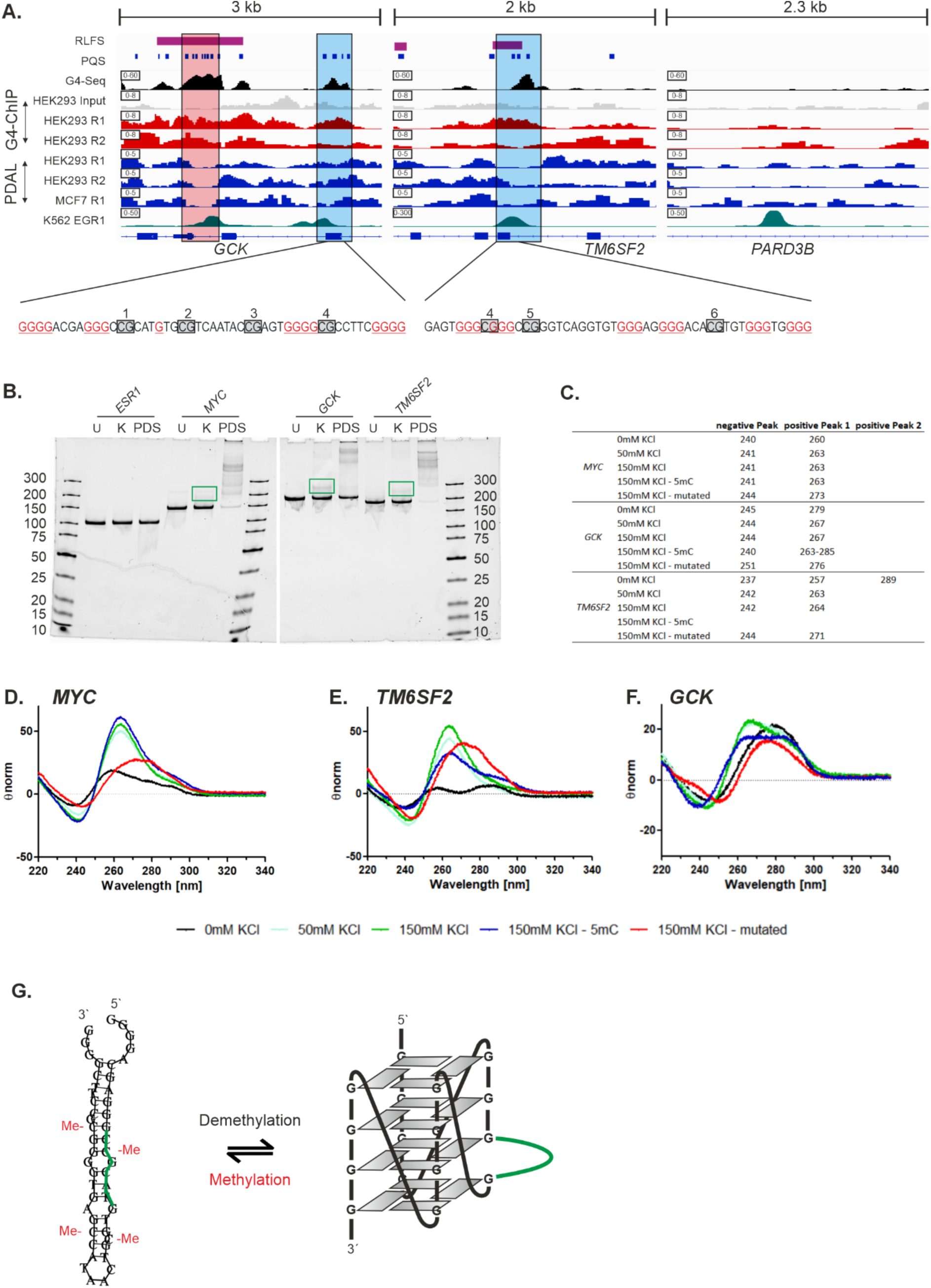
Non-B DNA formation in *GCK* and *TM6SF2*: **A.** Shows *GCK* (chr7: 44,184,901-44,187,919), *TM6SF2* (chr19: 19,380,358-19,382,362) and the non-G4 forming region in *PARD3B* (chr2:205,780,593-205,782,891). Data from different sources are shown in individual rows: Pink rectangles indicate R-loop forming sequences (RLFS) calculated by Quantitative Model of R-loop Forming Sequence (RLFS) finder (QmRLFS-finder)^76^. Blue rectangles represent putative G4 forming motifs (PQS) obtained from pqsfinder^77^, Black track: *in vitro* G4 formation stabilized with pyridostatin (PDS) in HEK293T cells^78^; Grey track: ChIP input control^79^; Red tracks: G4-ChIP-Seq tracks in HEK293 cells (2 replicates)^79^; Blue tracks: PDAL-Seq replicates 1 and 2 of HEK293 and MCF7 cells, Olive tracks: transcription factor EGR1 binding in K546 cells^82^. The sequences within *GCK* (antisense G4, sequence is given as reverse complement) and *TM6SF2* (sense G4) forming a G4 structure are detailed below. G-tracks are underlined, grey boxes indicate CpG positions. Blue highlighted areas: identified differentially methylated regions; light red highlighted area: a second regulatory region in *GCK,* which can form G4s. **B.** Native DNA PAA gels identify *in vitro* G4 formation in the G4 control region in *MYC*, *GCK* exon 7 and in the *TM6SF2* intron 2-exon 3 boundary, but not for *ESR1* (negative control). PCR products are loaded as unfolded (50 mM Tris-HCl pH 8.3) (U), folded with 150 mM KCl and heat induction (K), and folded with 150 mM KCl, heat induction and 10 µM PDS (PDS). **C.** positive and negative peak maxima obtained from CD spectra for *MYC*, *GCK* and *TM6SF2* with increasing K+ concentrations, for the methylated or mutated oligos. Individual CD spectra are shown for **D.** *MYC*, **E.** *TM6SF2* and **F.** *GCK*. **G.** Depicts the possible refolding of the methylated hairpin to a parallel G4 structure caused by demethylation at the *GCK* exon 7 region. The green loop indicates the bulge sequence which interrupts G-run two.

Next, we analyzed EGR1 ChIP-Seq data in K562 from ENCODE^82^. Unfortunately, there were no data available in HEK293 or MCF7 cells. Strong EGR1 signals were detected co-occurring with the G4-Seq peak and the footprinting valley of PDAL-Seq. Currently we do not know, if non-B DNAs are formed in K562 cells. The negative G4 region in *PARD3B* showed also a EGR1 binding site indicating that EGR1 does not only target G4 regions.

To independently evaluate the G4-forming ability of the target regions, G4 folding was confirmed by native DNA polyacrylamide (PAA) gel electrophoresis (Fig. 2B). The *GCK* and *TM6SF2* target regions and two controls, *MYC* as a positive G4-folding region and an enhancer region of *ESR1* as a G4-negative region, were PCR amplified and separated on a native 10% DNA PAA gel. Strikingly, both *GCK* and *TM6SF2*, as well as *MYC* showed significant shifted bands after the G4 folding procedure with potassium and heat induction (Fig. 2B, labeled with K), and even stronger after G4-stabilization with PDS (Fig. 2B, labeled with PDS), compared to their unfolded controls only containing Tris-buffer (Fig. 2B, labeled with U). This effect was not detected for the negative control region *ESR1*. In agreement with the native gel electrophoresis, circular dichroism (CD) spectroscopy experiments (Fig. 2C-F) experiments showed G4 formation for all regions enhanced by increasing K^+^ concentrations. *MYC* (Fig. 2C and D) and *TM6SF2* (Fig. 2C and E) showed characteristic parallel G4 structures with a negative peak at 240nm and a positive peak at 260nm. DNA methylation within *MYC* did not change the conformation of the G4 nor did it lead to a loss of the formed structure. For *GCK* we noticed an interesting pattern. While the increased K^+^ concentrations led to parallel G4 formation visible due to a shift in the positive peak from 279nm to 267nm, the methylated *GCK* oligo showed a broad positive peak from 263-285nm, which resembled a hairpin structure (Fig. 2D and F). This conformational change was probably caused by the methyl-groups which destabilize the G4, since there were six bases within the bulge followed by one G belonging to G-run two and 16 bases in loop two, which are able to pair (Fig. 2G). In all CD-spectra, the mutated oligos showed a clear shift of the positive peak to around 280nm, indicating the loss of G4 formation.

### Differentially methylated regions in *GCK* and *TM6SF2* harbor regulatory functions and contact neighboring genes

To assess the regulatory potential of the identified DMPs located in G4-forming regions in *GCK* and *TM6SF2*, we analyzed the activating H3K27ac, H3K4me1, H3K4me3 and the repressing H3K27me3 histone marks in H1-hESC (human embryonic stem cell line, red), HepG2 (hepatoblastoma cell line, blue), NHLF (lung fibroblasts, orange), K562 (lymphoblasts, brown), NHEK (normal human epidermal keratinocytes, violet), and MCF7 (breast cancer, green) cell lines in IGV from the ENCODE project^82^. In *GCK* exon 7 H3K4me1 activating histone marks in K562 cells and to a lesser extent also in HepG2 cells were found, whereas NHLF and NHEK cells harbored the repressive H3K27me3 mark (Fig. 3A, left panel). MCF7 cells showed H3K27ac marks near the target region in exon 7. Furthermore, the downstream regions overlapping exon 8-10 showed cell-type specific marks as indicated by H3K4me1 in HepG2 and K562 cells, but H3K27me3 in H1-hESC, NHLF and NHEK cells. For the *TM6SF2* intron 2-exon 3 boundary (Fig. 3A, right panel) we found strong signals for the activating H3K4me1 histone modification in K562 cells, but H3K27me3 signals in NHEK and NHLF cells. HepG2 cells showed H3K4me1 and H3K4me3 marks in the promoter region of *TM6SF2*. In conclusion, the analysis of ENCODE histone modifications revealed cell type specific regulatory elements for both target regions.

**Figure 3.**
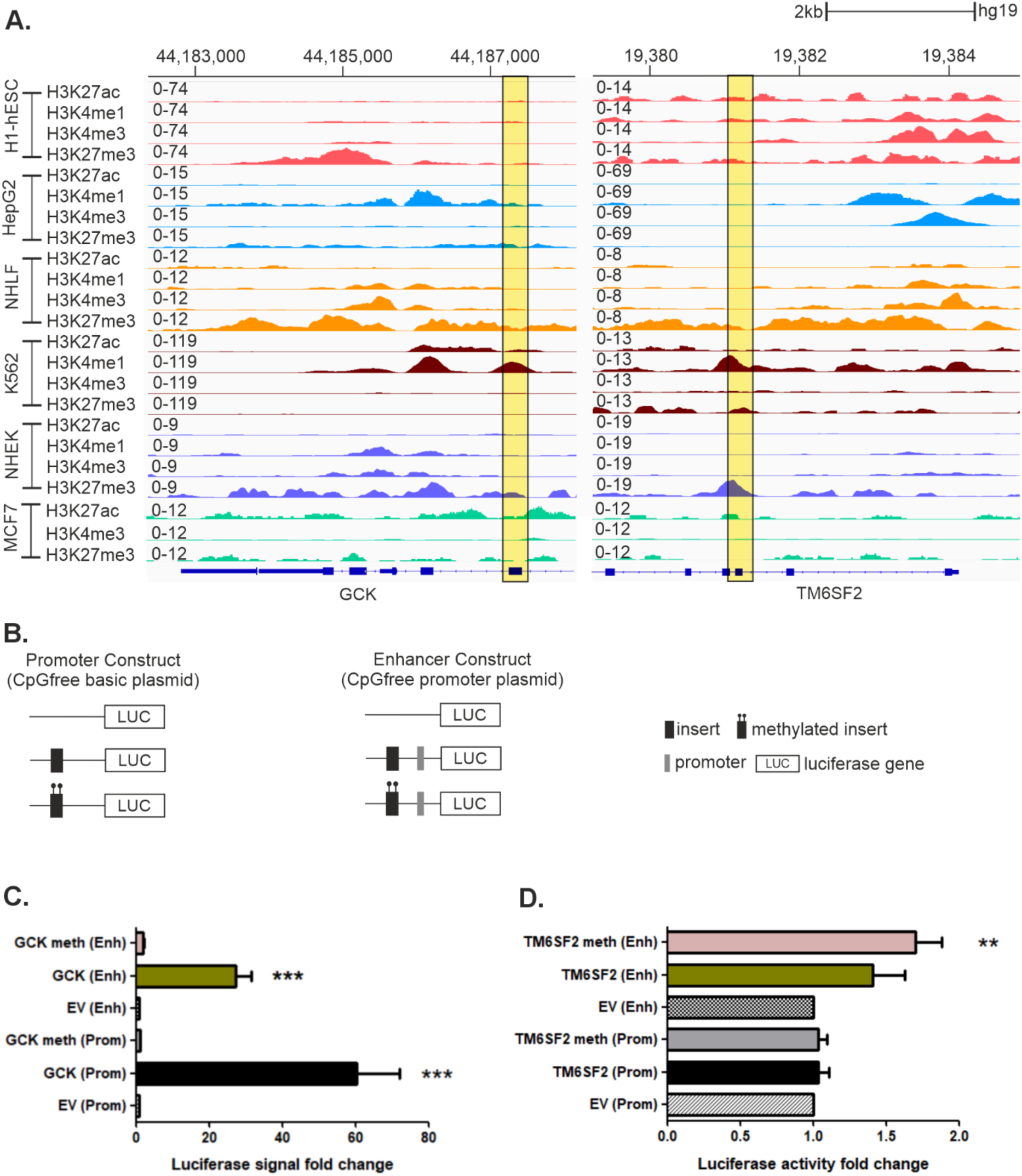
Regulatory roles of *GCK* and *TM6SF4* target regions. **A.** Activating (H3K27ac, H3K4me1 and H3K4me3) and repressing (H3K27me3) histone modifications obtained from ChIP-seq experiments from ENCODE for H1-hESC (stem cells, red), HepG2 (hepatoblastoma, blue), NHLF (healthy lung, orange), K562 (blood, brown), NHEK (normal human epidermal keratinocytes, violet) and MCF7 (breast cancer, green) cell lines ^82^ in *GCK* (left panel) and *TM6SF2* (right panel). For the hg19 genome assembly ENCODE did not provide H3K4me1 data for MCF7 cells. Yellow highlighted areas represent the pyrosequencing regions. Here we showed a cell-type specific pattern of activating and repressing histone marks. **B.** The CpG-free basic vector was used to test for a putative promoter function and the CpG-free promoter (containing a minimal promoter) vector to test for a putative enhancer function of the differentially methylated regions in a luciferase gene reporter assay in **C.** *GCK* exon 7 and **D.** in the *TM6SF2* intron 2-exon 3 boundary, in a methylated and unmethylated state compared to the empty vector (EV). Pairwise comparison against empty vectors (Welch two sample t-test), \**p*<.05, \*\**p*<.01, \*\*\**p*<.001.

To analyze the effect of the identified differentially methylated regions on gene expression, we performed *in vitro* luciferase reporter assays. Therefore, we cloned the regions of interest in two different CpG-free vectors upstream of the luciferase reporter gene and tested for their potential promoter and enhancer functions in a methylated and unmethylated state (Fig. 3B). We detected both, a significant promoter (60.3±11.3-fold, Welch two sample t-test, *p*=.0004,) and enhancer (27.4±3.9-fold, Welch two sample t-test, *p*=.0001, Fig. 3C) function for the *GCK* region compared to the empty vector, reflecting well the annotated ChromHMM function in HepG2 in the UCSC genome browser. However, when the same region was methylated prior to the analysis, both functions were lost. In contrast, we did not find a promoter function for the DMR located in *TM6SF2*, but the methylated construct showed a 1.7±0.2-fold (Welch Two Sample t-test *p*=.004, Fig. 3D) elevated enhancer function.

To predict the potential neighboring targets of the identified *GCK* and *TM6SF2* regulatory regions, we analyzed Hi-C data in HepG2 cells^82^ and from HCC patient material^83^ using the 3D Genome Browser^84^. *GCK* is located in a topological associated domain (TAD) of approximately 500 kb. Strikingly, within this TAD several genes are known for their association with metabolic diseases and cancer like AE binding protein 1 (*AEBP1*)^85,86^, YKT6 v-SNARE homolog (*YKT6*)^87,88^, upregulator of cell proliferation (*URGCP*)^89,90^ and phosphoglycerate mutase 2 (*PGAM2;* for more details see Supplement Section 4)^91^. When comparing the interaction frequencies of HepG2 cells and HCC decreased signal intensities were observed (Fig. 4A, comparison between TAD interaction frequency in HepG2-top and HCC-bottom). By analyzing the potential contact points of the *GCK* DMP, we noticed that this region stays in contact with an enhancer region in *GCK* intron 1 (∼chr7:44,157,293-44,158,172; hg38) and a proximal enhancer region between myosin light chain 7 (*MYL7*) and DNA polymerase delta 2 (*POLD2,* Fig. 4B). *TM6SF2* is located in a TAD together with UPF1 RNA helicase and ATPase (*UPF1*) and neurocan (*NCAN*, Fig. 4C), associated with MAFLD^92^ and HCC^93^. The interaction analysis showed that the *TM6SF2* DMP contacts a promoter-like region within intron 3 of hyaluronan and proteoglycan link protein 4 (*HPLN4*, ∼chr19:19,260,457, hg38) as well as intron 8 of SURP and G-patch domain containing 1 (*SUGP1*, ∼chr19:19,280,457, hg38), which was identified as regulator of cholesterol metabolism (Fig. 4D)^94^.

**Figure 4.**
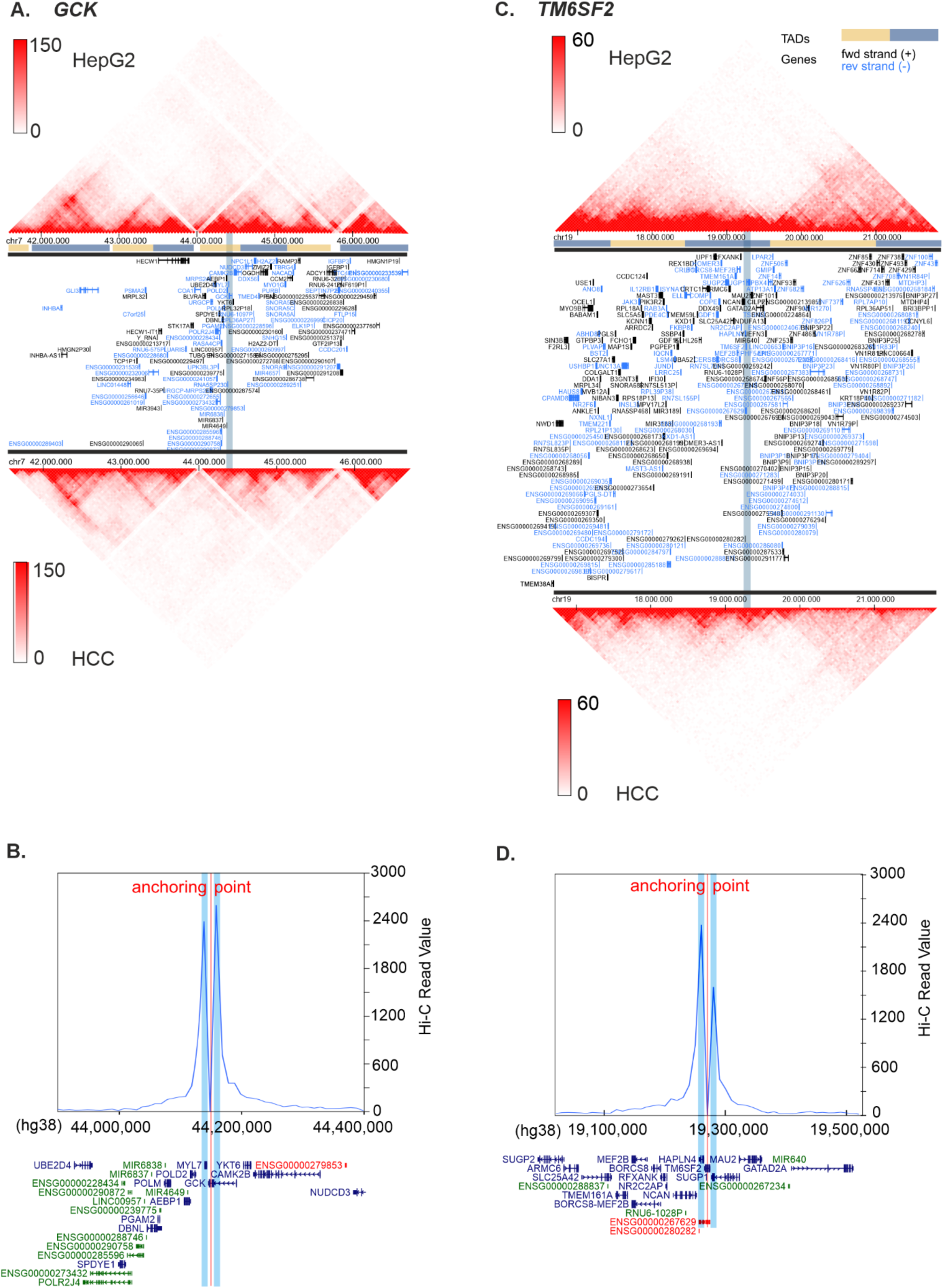
HiC data analysis in HepG2 cells and in HCC tumor material visualized in 3D genome browser^84^. **A.** TAD formation on chr. 7 around the *GCK* DMP in HepG2 (upper panel)^82^ and HCC tumor tissue (lower panel)^83^. **B.** Analysis of potential contact points of the regulatory region (red vertical line) shows interactions (blue highlighted areas) with *GCK* intron 1 and an intergenic region between *MYL7* and *POLD2* near an enhancer region. **C.** TAD formation on chr. 19 around *TM6SF2* in HepG2 (upper panel)^82^ and HCC tumor tissue (lower panel)^83^. **D.** Analysis of potential contact points of the regulatory region (red vertical line) shows interactions (blue highlighted areas) with *SUGP1* and *HAPLN4*. Chromosome annotations are given in the newer assembly hg38.

In summary, both *in silico* data for histone marks and HiC data, as well as the luciferase reporter assay provide evidence that *GCK* exon 7 and the *TM6SF2* intron 2-exon 3 boundary harbor important regulatory elements, which are not only related to metabolic diseases, but also could play roles in tumor formation and prognosis. Furthermore, our analysis points out that these regions contact neighboring genes and that changes in their methylation patterns potentially also affect the interactions with these elements. This strongly indicates that changes in DNA methylation at these regulatory elements are not only associated with MAFLD but could also be linked to the enhanced cancer risk in MetD patients corroborated also by the changes in the HiC data within these TADs.

## 3. Discussion

Metabolic diseases often arise due to a complex interplay of genetic and epigenetic processes. Whereas the genetic risk is predefined by the base composition, the epigenetic landscape is flexible and heavily influenced by the individualś environment. To assess epigenetic changes in two metabolic disease associated genes, *GCK* and *TM6SF2,* we analyzed 5mC in 148 patients grouped in five different categories: lean- and obese healthy subjects, lean and obese MAFLD, as well as patients suffering from advanced MetDs. We focused on two intragenic regions, annotated as regulatory regions in ChromHMM and predicted as non-B DNA forming sites (G4s and R-loops), which are not overlapping with CGIs. Significant 5mC changes were detected for both target genes. While *GCK* exon 7 was significantly hypomethylated especially at CpG 3 in all groups compared to lean healthy subjects (Fig. 1A), we observed a diverse pattern for the *TM6SF2* intron 2-exon 3 boundary. Lean- and obese MAFLD subjects showed hypomethylation, whereas hypermethylation was detected in advanced MetD (Fig. 1B). This pattern likely mirrors the elevated blood glucose levels observed also as HbA_1c_>7% within the advanced MetD group, potentially leading to hypermethylation. Furthermore, it also could indicate a change in blood cell type composition in this group, which also influences DNA methylation patterns^95^. Since increasing chronological age is one major risk factor for developing metabolic diseases (reviewed in^68^), our findings of a significant age correlation with DNA methylation within *GCK* and *TM6SF2* were expected. Nevertheless, after limiting the analysis to patients between 42-60-years of age, significant differential methylation was still detected for both genes, pointing towards a combined disease and aging mechanism. In addition, we were able to exclude an influence of the *TM6SF2* disease risk SNP rs58542926 C>T on DNA methylation in this region.

Moreover, we confirmed our results in hepatocellular carcinoma versus adjacent normal tissue obtained from *TCGA* data^71^. Although in contrast to cg11911689 (*TM6SF2,* Fig. 1D), differential methylation was not significant in cg23381646 (*GCK,* Fig. 1C), both CpGs showed a highly variable 5mC range, which is a sign for DNA methylation instability as previously observed, e.g. in Alzheimer’s disease^96^. Furthermore, DMPs in *TM6SF2* were hypermethylated in advanced MetD patients, whereas the TCGA data from HCC tissue material showed hypomethylation. The observation of opposing directions of differential methylation in blood versus the affected tissue is not unusual. In our latest epigenome-wide association study, a regulatory region upstream of NHL repeat containing 1 (*NHLRC1*) was found to be hypervariable, and hypermethylated in blood of lung cancer patients, but hypomethylated in lung cancer versus normal adjacent tissue ^97^. Although blood DNA methylation patterns are not necessarily reflecting the changes in the affected tissues, especially in metabolic diseases where blood glucose and triglyceride or cholesterol levels are constantly elevated when untreated, these patterns could direct us towards the affected target genes.

The analysis of gene expression in liver tissue of MAFLD and HCC patients demonstrated their differential expression. In line with the data from the liver expression database (Fig. 1E-G)^72^, Ilieva and colleagues found that *GCK* expression was increased by 2.48-fold (false discovery rate (FDR) adjusted *p*=1.08E-05) in MAFLD compared to obese-healthy subjects and by 1.83-fold (FDR adjusted *p*=1.21E-05) in advanced MetD patients compared to lean-healthy subjects in blood^98^. In contrast, *GCK* was downregulated in HCC compared to normal tissue from TCGA (Fig. 1E), which is at least in part explained by the replacement of *GCK* by hexokinase 2 in tumor cells^99^. The same trend was observed for *TM6SF2*, which was overexpressed in MAFLD, but decreased in HCC. Strikingly, the overlapping sense lncRNA ENSG00000267629 was overexpressed by 1.92-fold in the liver expression database in MAFLD compared to normal tissue^72^ and even by 3.19-fold (FDR adjusted *p*=.0005898) in MAFLD compared to obese-healthy subjects in the study of Ilieva et al.^98^. Furthermore, the expression level of this lncRNA was prognostic for the survival of intrahepatic cholangiocarcinoma (iCCA) with a hazard ratio (HR) of 1.582 (1.143-2.190, 95% CI, *p*=.006). Patients with a higher survival probability showed lower expression of this lncRNA transcript ^65^. Moreover, it was also prognostic for patients suffering from acute myeloid leukemia (AML)^64^.

Summarizing, DNA methylation of the target regions is not only detected in early stages of a metabolic disease (in *TM6SF2* after age correction even already in obesity) but also in HCC, indicating that DNA methylation changes observed in cancer are in some cases already detectable in a precancerous state which is in keeping with the increased risk for HCC observed in MAFLD patients.

G4s have emerged as major contributors to DNA methylation regulation and therefore, the detection of DMPs in G4 forming motifs is not new. Nevertheless, the shared occurrence of aberrant DNA methylation in an early disease stage such as obesity or metabolic diseases prior to cancer formation, implies an urgent need to understand their functions. We were able to show G4 formation in the *TM6SF2* intron 2-exon 3 boundary with PDAL-Seq in living cells (Fig. 2A), which adopts a parallel structure according to the CD-spectra (Fig. 2E). Overall, the DNA methylation in normal liver tissue (Fig. 2D) and also in several cell lines is low (Supplement Fig. 6A). Therefore, stable G4 formation is most likely occurring in this region. In contrast, *GCK* exon 7 harbors a ssDNA stretch, but slightly downstream of the G4-Seq peak. Since many cell lines such as HepG2 or MCF7 (Supplement Fig. 6B) as well as the normal liver tissue (Fig. 1C) are highly methylated in this region, stable G4 formation is unlikely, rather than a hairpin, as inferred from the CD spectra of the methylated *GCK* oligo. Hence, when demethylation occurs, a parallel structure is formed (Fig. 2F and G). This observation is in line with the structural change in *c-KIT* and *HRAS* caused by DNA methylation^52^. Furthermore, PDAL-Seq revealed the formation of G4 structures also exon 8-10, which has been shown as significantly hypomethylated in patients suffering from coronary heart disease (49.77 ± 6.43%) compared to controls (54.47 ± 7.65%, *p*=.018)^63^. CVDs are a common comorbidity of MAFLD and T2D and therefore this hypomethylation reflects the same pattern than the observed hypomethylation at exon 7 in our cohort.

When differential methylation occurs within regulatory elements, it inevitably affects gene expression patterns. Both regions exhibit cell-type-specific histone marks (Fig. 3A), luciferase reporter assays demonstrate that their regulatory functions are activated in a DNA methylation-dependent manner (Fig. 3B and C), and HiC data indicate that these elements interact with neighboring genes (Fig. 4). Notably, DNA methylation of both regions is highly variable, and exhibit enriched binding for the zinc finger transcription factor EGR1. While it has been reported that G4s drive DNA methylation processes by sequestering DNMT1 to the G4 and cause its inactivation^26^, we propose a slightly different, DNMT1-independent mechanism for these regions.

EGR1 recognizes a G-rich motif^100^ and is predicted as G4 binding protein^101^. It recruits the ten eleven translocase 1 (TET1), leading to DNA demethylation processes adjacent to its binding sites^102,103^. Its binding is likely not restricted solely to folded G4s, as the *PARD3B* region, which lacks any non-B DNA structure, also shows EGR1 binding. EGR1 has been shown as important transcription factor regulating inflammatory genes by binding to their enhancers^104^. EGR1 is upregulated in MAFLD, and a its knockdown resulted in a reduced expression of pro-inflammatory genes in hepatocytes^105^.

Given that the *GCK* exon 7 G4-CpGs are methylated in normal liver tissue, as well as different cell lines, we hypothesize that local demethylation processes initiated by EGR1 binding, facilitate G4 formation, thereby allowing transcription factor binding and activation of the regulatory element (Fig. 5A). This is supported by experiments showing a ∼60-fold increase in promoter activity and a ∼27-fold increase in enhancer function when GCK exon 7 is unmethylated (Fig. 3C). The *TM6SF2* intron 2-exon 3 boundary shows clear non-B DNA formation, and methylation reduces the stability of its G4 structure (Fig. 2E). DNA methylation at the nearest CpG position is lower in the target region in *TM6SF2* than in *GCK* in both normal tissues and cell lines (Fig. 1D, Supplement Fig. 6), implying that this element is active in a healthy state, at least in a cell-type specific manner. The strong binding of EGR1 to the region suggests, that DNA demethylation is facilitated by EGR1 and TET1 recruitment and a stable parallel G4 is formed. However, increased DNA methylation in this region, as seen in advanced MetD patients or in cases of high methylation instability as observed in HCC, could impact the regulatory element’s activity. Notably, we analyzed only a rather small part of the UCSC-annotated regulatory element, which may have caused the overall low detected enhancer strength. In line with our study, but investigating long G4-forming regions, Williams et al. found that G4-forming regions harbor promoter and enhancer functions and that the most prominently enriched interacting transcription factor binding site was that of EGR1^32^.

**Figure 5:**
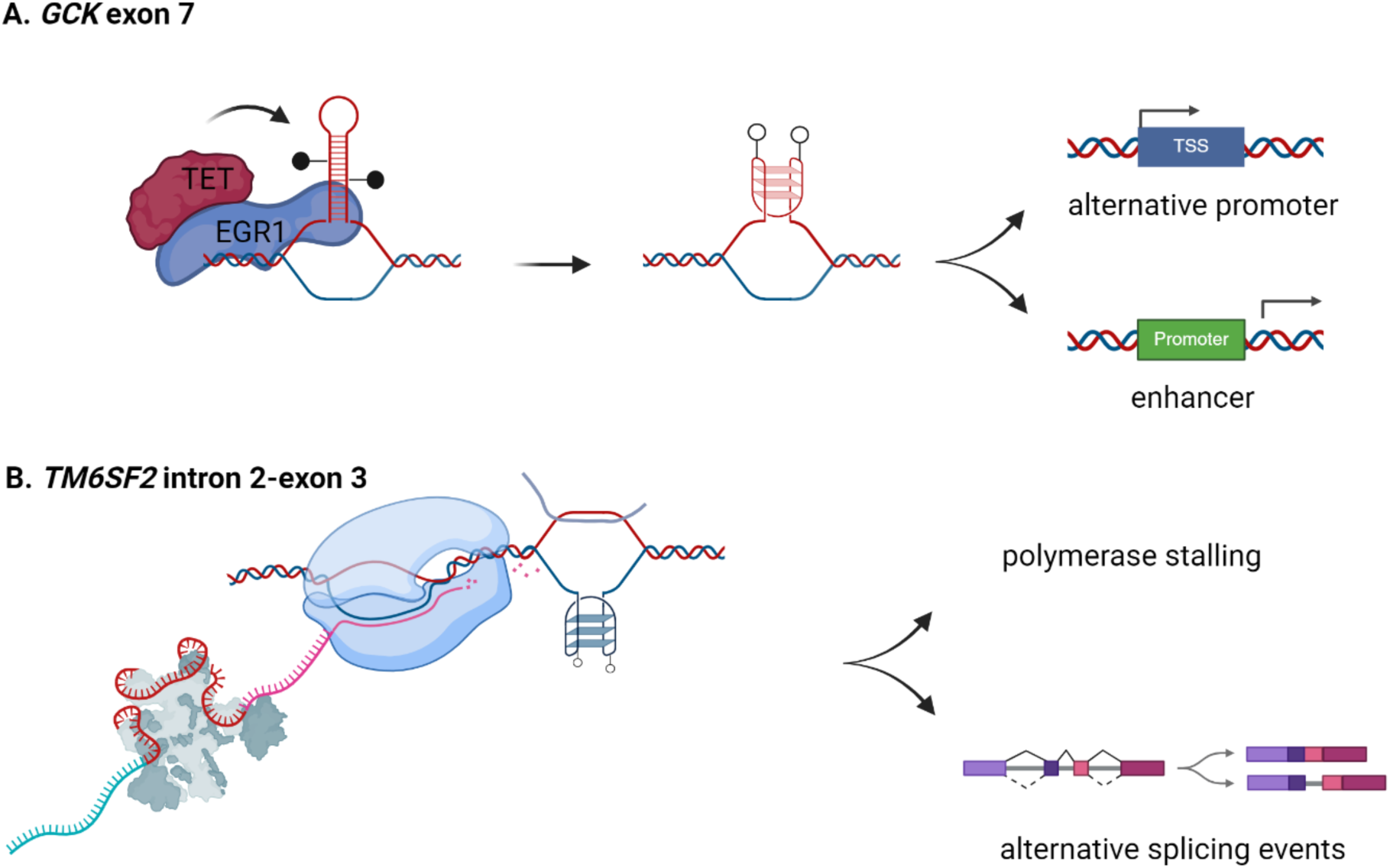
Mechanistic explanation. **A.** Hypomethylation of *GCK* exon 7 leads to the formation of a G4 structure which in turn activates the alternative regulatory element. **B.** Methylation instability at the cassette exon and G4 forming region in the *TM6SF2* intron 2-exon 3 boundary could cause polymerase slow down or stalling and alternative splicing events leading to a readthrough of *TM6SF2* to *HAPLN4*, forming the long non-coding RNA. Created with BioRender.com.

Alternative promoters play critical roles in hepatocellular and breast cancer, two cancer types highly linked to metabolic diseases and obesity^10,12^. Genes involved in glycolysis and cholesterol metabolism show elevated expression from alternative promoters, which typically exhibit low H3K4me3 levels and hypomethylation in CpGs located in non-CGI promoters^58^. CGIs are underrepresented in alternative promoters, and Nepal et al. propose that activation of these promoters in HCC occurs through demethylation of CpG-poor intragenic regions^58^, similar to the investigated G4-forming regions.

Additionally, the TM6SF2 intron 2-exon 3 boundary could be subject to alternative splicing, since exon 3 of *TM6SF2* is annotated as a cassette exon, and G4s are enriched at distal splice sites of such exons^33^. Splicing factors like U2AF1, which recognizes the 3ʹ splice site, are among the top candidates associated with G4s^37^. G4 formation regulates alternative splicing of BCL-x, since stabilization of the G4 structure promotes the expression of the pro-apoptotic BCL-xS isoform and triggering apoptosis^106^. However, this requires critical discussion since the G4 motif is found on the template strand, meaning the pre-mRNA cannot not form the G4. Probably the G4 creates an obstacle for the RNA polymerase, causing reduced polymerase speed or stalling. Of note, proteins recognizing methylated DNA can slow down RNA polymerase II, thereby indirectly influencing alternative splicing mechanisms^107^. In line with this, targeted demethylation via TET1 linked to dCas9 causes exon skipping^108^. Aberrant splicing regulation can result in readthroughs to neighboring genes during stress conditions^109^, which could explain the overexpressed lncRNA ENSG00000267629 in MAFLD. In summary, the observed differential methylation in MAFLD groups in HCC, in combination with G4 formation, suggests a rather complex interplay between transcription and probably also alternative splicing processes (Fig. 5B).

### Conclusion

We were able to show aberrant DNA methylation within the investigated intragenic regions of *GCK* and *TM6SF2* in metabolic diseases as well as in HCC and that these structures are alternative regulatory elements potentially controlled by G4 formation. The link between G4s and cancer is well known. While the number of G4s naturally decreases during cell differentiation^110^, G4s are increased in tumor tissue^111^ and in the human keratinocytes (HaCaT) cancer cell line versus normal human epidermal keratinocytes NHEK^112^. Even the cell-type specific transcriptome has been shown to be shaped by G4s^113^. Therefore, G4s harbor a great potential as novel therapeutics and there are already G4 ligands which are used in clinical trial for solid tumors with mutations in homologous repair genes or pancreatic cancers (reviewed in^114^). Recently it has been shown that pathologically high glucose concentrations cause the formation of a stable promoter-G4 and in turn result in significantly increased gene expression compared to normal glucose concentrations^45^. We suggest that a similar mechanism also applies for MetD patients since they are exposed constantly to high glucose and high fatty acid concentrations, if not treated properly.

Although further research is necessary to get a deeper understanding of the mechanisms involved, we were able to connect detected DMPs in metabolic risk genes to G-quadruplex formation in a patient cohort suffering from metabolism dysfunction-associated fatty liver disease and type II diabetes, for the very first time. Unfortunately, we are currently lacking G4 formation data in human liver biopsies, since this type of material is rare. Although the assessment of anthropometric and blood parameters, abdominal MRI and liver biopsies are the current standard for MAFLD diagnosis, liver biopsies cannot be applied on a broad scale in asymptomatic obese or lean-MAFLD subjects for ethical reasons or low patient acceptance. In conclusion, since metabolic diseases are linked to an elevated cancer risk, and aberrant DNA methylation as well as elevated non-B DNA formation occurs during cancer formation, these mechanisms could point to a common mechanism of metabolic diseases and liver cancer development. Understanding this intricate connection especially in intragenic regions with alternative regulatory roles is paramount in unraveling the complexities of metabolic diseases, potentially leading to G4s as novel targets for metabolic disease treatment as already evaluated for solid cancers.

## 4. Methods

### Study design

We designed a nested cohort study by selecting 148 blood samples from Caucasian origin collected at the University Hospital Salzburg, Austria, which fit into one of five groups according to the criteria below. The local ethics committee (Ethikkommission des Landes Salzburg No. 415-E/1521/6-2012, and an ethics approval by the Paracelsus Medical University Salzburg No. 01-2022) approved the study and informed consent was obtained from all participants. The study was conducted in accordance with the guidelines of the World Medical Association’s Declaration of Helsinki.

Alanine transaminase (ALT) and aspartate transaminase (AST) levels ≤40 U/L for women and ≤50 U/L for men, with or without increased levels of γ-glutamyl transpeptidase (GGT), were considered as normal liver tests. Routine clinical parameters were determined during sampling at the outpatient clinics of the University Hospital Salzburg, Austria.

(1) Lean healthy subjects (LH, n=24): body mass index (BMI) ≤25.0 kg/m^2^, normal liver tests, normal ultrasound and no components of MetS according to ATPIII criteria^115^
(2) Obese healthy subjects (OH, n=31): metabolic healthy obese (MHO) BMI ≥30.0 kg/m^2^, normal ultrasound and normal liver tests, and ≤2 components of MetS (obesity and one additional metabolic syndrome defining criterion allowed and glycated hemoglobin (HbA_1c_)<6.0%;
(3) Lean MAFLD subjects (LM, n=27): metabolic unhealthy normal weight (MUNW), BMI ≤25.0 kg/m^2^, ultrasound evidence of a fatty liver, with or without elevated liver tests, HbA_1c_<7%;
(4) Obese MAFLD subjects (OM, n=32): BMIs ≥30.0 kg/m^2^, unequivocal fatty liver ultrasound, with or without elevated liver tests, HbA_1c_<7%;
(5) Advanced MetD (aMetD, n=34): Elevated fibrosis-4 (FIB4) score>2.67 and/or established T2D with elevated HbA_1c_>7%.

### Sequence specific DNA methylation analysis by pyrosequencing

DNA was extracted from 200 µL whole blood with the ReliaPrep Blood gDNA Miniprep System (Promega, Austria) according to the manufactureŕs guidelines. Bisulfite (BS) conversion was performed with 500 ng of genomic DNA (gDNA) input material using a Bisulfite Conversion Kit (Zymo Research, Austria). DNA was eluted with 60 µL M-Elution buffer (∼8 ng/µL). PCR was performed in 25 µL reaction volume containing 200 µM dNTPs, 1 mM MgCl_2_, each 0.2 µM forward and reverse primer, 0.63 U HotStar Taq polymerase (Qiagen, Austria) and 16 ng (*P2RXL1*, SCGN and *NHLRC1*) or 32 ng (*EDARADD*, *IPO8*, *GCK* and *TM6SF2*) of BS-converted DNA to reach sufficient material for pyrosequencing. The PCR protocol included an initial heating step of 95°C for 15 minutes, followed by 45 cycles of 95°C for 30 seconds, 55-64°C primer annealing depending on the target (see Supplement Material & Methods section 1.1) for 30 seconds and 72°C for 15 seconds, as well as a final elongation step at 72°C for 7 minutes. Pyrosequencing was performed using 20 µL PCR product and 7.5 µM (*IPO8*, *GCK*, *TM6SF2, P2RXL1*, and *NHLRC1*) or 9 µM (*SCGN* and *EDARADD*) sequencing primer. Detailed assay design, primer sequences, annealing temperatures and sequencing conditions can be found in the Supplement (Material & Methods section 1.1). Data analysis was performed with Pyromark Q24 Advanced software, R version 4.3.1^116^ and R studio^117^.

### rs58542926 C>T single nucleotide polymorphism (SNP) genotyping by pyrosequencing

To infer the genotype of the rs58542926 C>T (E167K) risk variant, we amplified the region in 25 µL reaction volume using 200 µM dNTPs, 1 mM MgCl_2_, each 0.2 µM forward and reverse primer, 0.63 U HotStar Taq polymerase (Qiagen, Austria) and 10 ng genomic DNA. The PCR protocol included an initial heating step of 95°C for 15 minutes, followed by 45 cycles of 95°C for 30 seconds, 63°C for 30 seconds and 72°C for 15 seconds, as well as a final elongation step at 72°C for 7 minutes. For the pyrosequencing reaction, we used 20 µl PCR product and 7.5 µM sequencing primer. Detailed information is given in the Supplement Material & Methods section 1.1. Data analysis was performed with Pyromark Q24 Advanced software, R version 4.3.1 ^116^ and R studio ^117^.

### Gene expression data

Expression data for MAFLD and HCC were extracted from the liver expression portal (GepLiver)^72^. For the analysis, we downloaded the datasets for Bulk-01 (normal liver), Bulk-05-09 (including MAFLD) and Bulk-19 (TCGA; including HCC) datasets as listed in the GepLiver portal. All statistical analyses were performed using R version 4.3.1^116^ and R Studio^117^.

### Non-B DNA prediction tools

To predict non-B forming sites, we utilized pqsfinder for G4 forming sites^77^, QmRLFS-finder for R-loop forming sequences (RLFS)^76^, and the *RNAfold* web server of the University of Vienna/Austria for the folding of hairpins^118^.

### Non-B DNA detection with permanganate/S1 nuclease footprinting with direct adapter ligation and sequencing (PDAL-Seq)

PDAL-Seq was performed according to the protocol described in^81^. Further details can be found in the Supplement (Material & Methods Section 1.3). In short, 5-7 x 10^6^ cells were washed two times with 1x PBS without Ca^2+^ and Mg^2+^ and incubated in 1 mL low salt buffer (15 mM Tris-HCl pH 7.5, 15 mM NaCl, 60 mM KCl, 300 mM sucrose, 0.5 mM EGTA) for 1 min. 670 µL of freshly prepared KMnO_4_ stock solution (40 mM for HEK293, 70mM for MCF7) were added and incubated at 37°C in the dark for 80 sec. Subsequently, the reaction was stopped with 2.67 mL stop solution (50 mM EDTA, 700 mM β-mercaptoethanol, 1% SDS) and 79.5 µL proteinase K (20 mg/ml stock solution, Promega, Austria) were added. After overnight incubation at 37°C, DNA was extracted with phenol chloroform isoamylalcohol (PCI, Merck, Austria) and precipitated with 2 M NH_4_OAc and 100% ethanol. After resuspending the pellet in 1 mL of 10 mM Tris HCl pH 8.0 buffer, RNA was digested by adding 2.5 µL of 20 mg/mL RNase A (Promega, Austria) for 1 h followed by another PCI extraction. Free DNA ends were blocked with 500 U Terminal Transferase (Roche, Austria), 120 µM cordycepin-5’-triphosphate and 5 mM CoCl_2_ in 500 µL total reaction volume, followed by a PCI extraction and two ethanol precipitations to avoid cordycepin carry over. 40 µg DNA were digested with 20 U S1 nuclease (0.5 U/1 µg DNA, Promega, Austria) in a total reaction volume of 600 µL at 37°C for 20 min. The reaction was stopped by adding 10 mM EDTA. After PCI extraction, end repair using T4 DNA polymerase (NEB, Austria) was performed for 15 min at 12°C, followed by PCI extraction. 500 nM biotinylated P5 adapter (Merck, Germany) were linked with 2000 U T4 DNA ligase (Promega, Austria) in a final volume of 100 µL at 16°C overnight. To remove unligated P5 adapters, Agencourt SPRI bead cleanup (Beckman Coulter, Austria) was performed with 0.9 volumes of beads and eluted in 132 µL 1x TE buffer (10 mM Tris HCl pH 8.0, 1mM EDTA). Sonication was performed with an Covaris M220 focused-ultrasonicator in 130 µL microTube-AFA Fiber Pre-Slit Snap Cap tubes with the parameters: peak incidence power: 75, duty factor: 10, cycles per burst: 200 and treatment time: 80 sec, to reach a target peak of approximately 400 bp. Fragments were bound to streptavidin coated magnetic beads (Dynabeads^®^ M-280 kilobaseBINDER kit, Thermo Scientific, Austria) and carefully washed with 10 mM Tris-HCl pH 7.5. The beads were then resuspended in 100 µL end repair mix (NEBNext End Repair Module, NEB, Austria) and incubated at 20°C for 30 min. After washing with 10 mM Tris-HCl pH 7.5, P7 adapter ligation (1 µM adapters, 400 U T4 DNA ligase) was performed on a wheel tube rotator at 20°C for 4 h. Beads were washed with 10 mM Tris-HCl pH 7.5 and resuspended in 100 µL PCR mix and PCR was performed. Detailed information about PCR reaction and primers are provided in the methods paper^81^ and in the Supplement (Material and Methods section 1.3). SPRI bead purification removed unbound adapters and primer dimers and the quality of the resulting libraries was checked using an Agilent Fragment Analyzer. Library concentration was determined with a KAPA Library Quantification Kit (Roche, Austria) and sequenced on a NovaSeq platform with approximately 80 million reads. After demultiplexing the raw reads and quality control, reads were aligned to the human reference genome hg19 with the BWA-MEM aligner^119^. Subsequent filtering removed low-quality reads (mapping quality MQ <30) and ENCODE blacklisted regions^82^. BamCoverage files were generated with a 50 bp bin size and normalized to 1× coverage and visualized in IGV^120^. Data analysis was performed using the open-source platform Galaxy^121^.

### Studying *in vitro* G4 formation with native DNA polyacrylamide gels and circular dichroism (CD) spectroscopy

Target regions harboring the DMPs in *GCK* and *TM6SF2*, a positive control region in *MYC* containing a predicted G4 motif^122^, and a negative control region within a regulatory region of the estrogen receptor substrate 1 (*ESR1)* without a predicted G4 motif^123^ were PCR-amplified. To do so, we prepared a 10 µL PCR reaction containing 200 µM dNTPs, 1 mM MgCl_2_, each 0.2 µM forward and reverse primer, 0.25 U HotStar Taq polymerase (Qiagen, Austria) and 10 ng genomic DNA. The PCR protocol included an initial heating step of 95°C for 15 minutes, followed by 45 cycles of 95°C for 30 seconds, 54-63°C primer annealing depending on the gene for 30 seconds and 72°C for 15 seconds, as well as a final elongation step at 72°C for 7 minutes. The PCR fragments were cleaned-up with ReliaPrep DNA concentration kit (Promega, Austria) and eluted with dH_2_O. The amplicons were then diluted to a final concentration of 20 ng/µL in 50 mM Tris-HCl (pH 8.3). To enhance G4 folding, we prepared the same dilutions and added 150 mM KCl or 150 mM KCl and 10 µM pyridostatin (PDS, Merck, Austria) to the Tris-HCl buffer. The fragments were then heated to 95°C and cooled gradually down to 25°C in a thermocycler overnight. DNA fragments were loaded onto a native 10% DNA polyacrylamide (PAA) gel using 1× TBE buffer at 68 V (8 V/cm^2^) for 2h. DNA PAA gels were post-stained with GelRed (Biotium, VWR, Austria) for 5 min and visualized with a ChemiDoc station (BioRad, Austria). Detailed information can be found in the Supplement (Material & Methods Section 1.3) as well as the raw gel image in Supplement Fig. 7.

Circular dichroism (CD) spectra were obtained from single stranded DNA oligos from the predicted G4 motif as well as in a methylated and mutated form (Supplement Material & Methods Section 1.4). All oligos were reconstituted in 50 mM Tris-HCl (pH 7.5) and ranging from 0 mM, 50 mM to 150 mM KCl in a final volume of 200 µL with an oligo concentration of 5µM. CD spectra were recorded using the Chirascan™ Plus CD Spectrophotometer instrument (Applied Photophysics, United Kingdom) in a 0.5 cm path cuvette at 25°C. Spectra were scanned from 340 to 220 nm., with a step size of 0.2 nm and a bandwidth of 1 nm. Four spectra per sample were averaged. After blank correction, the CD ellipticity θ in millidegrees was normalized using the relation θnorm = θ / (32980 × c × l), where c is the DNA concentration in mol/L and l is the path length in cm.

### TCGA data and ENCODE histone mark and TF binding data analysis

DNA methylation and gene expression data were obtained from tumor versus normal tissue from patients suffering from hepatocellular carcinoma (HCC) via the Shiny Methylation Analysis Resource Tool (SMART), an interactive tool for analyzing *The Cancer Genome Atlas* (TCGA) data^124^.

*The Encyclopedia of DNA Elements* (ENCODE) data for H3K27ac, H3K4me1 and H3K4me3 from the liver hepatoblastoma cell line HepG2 and the lymphoblast cell line K562^82^ were downloaded and visualized using the Integrative Genome Viewer (IGV)^120^. Early growth response 1 (EGR1) chromatin immunoprecipitation and sequencing (ChIP-Seq) data from K562 cells were obtained from the ENCODE database^82^.

### Analyzing contact points in HiC datasets

Genome organization and chromatin interaction data (HiC) were obtained from HepG2 cells via ENCODE^82^, and for HCC material^83^, and visualized with 3D Genome Browser^84^. To detect potential interactions with surrounding genes, we used the coordinates chr7: 44,187,355 in hg19 (chr7:44,147,756; hg38) for *GCK* exon 7 and chr19:19,381,266 in hg19 (chr19:19,270,457; hg38) for *TM6SF2* intron 2-exon 3 for virtual 4C analysis using HiC data in the 3D Genome Browser^84^.

### Testing for regulatory functions using the luciferase reporter assay

To verify the regulatory role of the DMPs identified in *GCK* exon 7 and *TM6SF2* intron 2-exon 3 boundary, the regions were cloned into pCpG-free vectors (Invivogen, Belgium). The pCpG-free basic vector was used to detect potential promoter activity and the pCpG-free promoter vector contains a minimal promoter to test for enhancer activity of the regions of interest. Since these vectors do not contain any CpG dinucleotides in the backbone, *in vitro* DNA methylation can be performed with the *M.SssI* enzyme, which then selectively methylates only the inserted human *GCK* or *TM6SF2* region and allows to selectively study the effect of 5mC in the insert on luciferase gene expression.

Briefly, DNA was isolated from HepG2 cells and the regions were amplified in 50 µL reaction volumes containing 1x GC buffer, 200 µM dNTPs, 0.5 µM forward and reverse primer, and 0.5 U Phusion HSII polymerase (Thermo Scientific, Austria). The PCR protocol contained an initial heating step of 98°C for 30 seconds, followed by 45 cycles of 98°C for 10 seconds, 69°C for 10 seconds and 72°C for 15 seconds, as well as a final elongation step at 72°C for 5 minutes. The PCR primers contained restriction enzyme (RE) cutting sites for *NsiI* (NEB, Austria) and *BamHI*/*HindIII* (NEB, Austria) at the 5’ends of the forward and reverse primers, respectively. 500 ng of each, PCR amplicon and plasmid, were double-digested at 37°C for 20 min and cleaned up with a DNA cleanup and concentration system (Promega, Austria). Detailed information can be found in the Supplement (Material & Methods section 1.5).

The ligation reaction was performed in a 1:5 plasmid to insert copy number ratio using T4 DNA ligase (NEB) at 16°C overnight. 5 µL of the ligated construct were transferred to chemically competent *E.coli* GT115 cells (*pir* mutant strain, Invivogen, Belgium) and spread on lysogeny broth (LB) plates containing zeocin (50 µL/ml; Invivogen, Belgium). Singe colonies were picked and incubated in 3 ml LB-zeocin medium overnight. Plasmid was extracted with PureYield Plasmid Midiprep System (Promega, Austria) and successful cloning was verified by Sanger Sequencing at LGC genomics (Germany). 1 µg of plasmid was incubated with 8 U *M.SssI* (NEB) at 37°C for 4 h for *in vitro* DNA methylation. The reaction was heat deactivated at 65°C for 20 min, followed by plasmid cleanup with ReliaPrep DNA Clean-Up and Concentration kit (Promega, Austria).

24 h prior to transfection, 0.15 x 10^5^ HepG2 cells per well were seeded in 24 well plates with DMEM low glucose medium + 10% fetal bovine serum without antibiotics. Lipofectamin 3000 (Thermo Scientific, Austria) was used to transfect 500 ng plasmid (renilla luciferase) and 25 ng pGL4 SV40 background plasmid (firefly luciferase) and incubated overnight. After cell lysis, chemiluminescent read-out was detected on a Tecan Spark plate reader (Tecan, Austria). Renilla readouts were normalized to the firefly luciferase and finally, data were expressed as fold-change normalized against the empty vector.

### Statistical analyses

All statistical analyses were performed using R version 4.3.1 ^116^ and R Studio ^117^. Significant changes in DNA methylation in all groups were detected using analysis of variance (ANOVA) with *p*<.05. For individual group comparisons, Student’s t-test (*p*<.05) was applied. DNA methylation data from TCGA and gene expression data from GepLiver database were analyzed with Wilcoxon rank sum test using Benjamini Hochberg (BH) correction. Welch’s two sample t-tests were performed to calculate significant differences in luciferase reporter assays (*p*<.05).

## Data Availability

PDAL-Seq data are available at the Sequence Read Archive (SRA) under the BioProject number PRJNA1088738 (https://www.ncbi.nlm.nih.gov/Traces/study/?acc=PRJNA1088738&o=acc_s%3Aa).

## Author contribution

A. L. established PDAL Seq, designed assays and conceptualized the study; A. L., V. E., A. O., Z. L. and E. S. performed experiments, B. P. and E. A. collected and grouped the samples and gave substantial advice and intellectual input to the study; A. L. wrote the manuscript, A. L., E.A. and A.R. revised and edited the manuscript. All authors read and approved the final manuscript.

## Acknowledgments

We want to thank the Austrian Diabetes Association (ÖDG) very warmly for their support as well as all patients who contributed their blood samples to the study. Furthermore, we thank Kateryna Makova and Matthias Weissensteiner for plenty of fruitful discussions on PDAL-Seq and their input on G-quadruplex structures.

## Abbreviations

5mC: 5-methylcytosine
BS: Bisulfite conversion
ChIP-Seq: Chromatin immunoprecipitation and sequencing
DMP: Differentially methylated position
DMR: Differentially methylated region
G4: G-quadruplex
GCK: Glucokinase
HiC/4C: Chromatin conformation capture
MAFLD: Metabolism dysfunction-associated fatty liver disease MetD Metabolic disease
MetS: Metabolic syndrome
Non-B DNA: Non-canonical DNA structure
PDAL-Seq: Permanganate/S1 nuclease footprinting with direct adapter ligation
PQS: Putative G4 forming sequence
R-loop: RNA:DNA hybrid
RLFS: R-loop forming sequence
T2D: Type 2 diabetes mellitus
TF: Transcription factor
TM6SF2: Transmembrane 6 superfamily 2

