## Supplement for "G-quadruplex forming regions in *GCK* and *TM6SF2* are targets for differential DNA methylation in metabolic disease and hepatocellular carcinoma patients"

University of Salzburg
Hellbrunnerstraße 34
5020 Salzburg
Austria

### 1. Supplementary Material and Methods

#### 1.1. Pyrosequencing

##### 1.1.1. PCR and Sequencing Primers

| Gene | CG ID/region | Primer | Primer Sequence 5'-3' | Amplicon length [bp] | $T_m$ [°C] |
|---|---|---|---|---|---|
| ***P2RXL1*** | cg05442902 | Forward | AGTTAGATAAGTAGATAAAGATGGGGATGG | 106 | 60 |
| | | Reverse-biotinylated | ACCCTCTCACTCTATACCTCTTAA | | |
| | | Forward-Sequencing | GGTTTTTGGTGTAGGG | | |
| ***SCGN*** | cg06493994 | Forward | TGAGTTTGGAGTTAAGTTTATATTTGAGTG | 273 | 62 |
| | | Reverse-biotinylated | CCTCCCCAACAACAATTACTC | | |
| | | Forward-Sequencing | GGTGTTGAAGAAGAAGATTAGGTAT | | |
| ***EDARADD*** | cg0980672 | Forward | TTTGGAGTTTGTTATGGAAGAAGT | 124 | 58 |
| | | Reverse-biotinylated | AAAAAAATTTATCCTCCCACCTACA | | |
| | | Forward-Sequencing | TGTTATGGAAGAAGTAATAGAT | | |
| ***IPO8*** | cg19722847 | Forward | TGGTAGGGAGTTTGTTTATAGTTGT | 119 | 58 |
| | | Reverse-biotinylated | ACAAATCTATCTTAAAACAAACCCAATTAC | | |
| | | Forward-Sequencing | GGATGGTGTGATATTTTTT | | |
| ***NHLRC1*** | cg22736354 | Forward-biotinylated | AATGGGTATTAGAGGGTTAGAGT | 150 | 55 |
| | | Reverse | AATCAACCTACTCCAATACAAAATATACTT | | |
| | | Reverse-Sequencing | AAAATATACTTTAAAAAATTTAACC | | |
| ***TM6SF2*** | Intron 2-exon 3 | Forward | GGAGGTAGTGAATAGTGTTATTTTAGTA | 400 | 60 |
| | | Reverse-biotinylated | ATAATCTCATCCCTAACCTTAACTTCAA | | |
| | | Forward-sequencing | GAAGAGAGATGATGAGGTTTATA | | |
| ***GCK*** | Exon 7 | Forward | GTTATATGGAGGAGATGTAGAATGT | 216 | 58 |
| | | Reverse-biotinylated | ACACCAAAACCACCCAAAACCTAAATTATA | | |
| | | Forward-sequencing | AATGTGGAGTTGGT | | |
| ***TM6SF2*** | rs585429 C/T | Forward-biotinylated | TAGGGGATGGTGAGGAAGAAGG | 60 | 63 |
| | | Reverse | CACCATGGAAGGCAAATACAGCT | | |
| | | Reverse-sequencing | GAAGGCAAATACAGCT | | |

##### 1.1.2. Pyrosequencing Assays/Sequence to Analyze

| Gene | Sequence before BT conversion | Sequence after BT conversion | Sequencing Primer conc [µM] |
|---|---|---|---|
| P2RXL1 | AGCAGCGTATGCAAAGA | GGAGGGGGACGAGGGCCGCATGTGCGTCAATACCGAG TGGGG | 7.5 |
| SCGN | CGGCGCGGAAGGACCTAGA | YGGYGYGGAAGGATTTAGA | 9 |
| EDARADD | TGCGAGAAGATGCTCGCTG | TGYGAGAAGATGTTYGTTG | 9 |
| IPO8 | GTTAGTCCGAGAACTGT | GTTAGTTYGAGAATTGT | 7.5 |
| NHLRC1 | ACCGGCAGCAGCGGCGCCCGCG | ATYGGTAGTAGYGGYGTTYGYG | 7.5 |
| TM6SF2 | ACCGAGGTGAAGGCGAAGACAGCGAAGACTGCAGTGAG TGGGCGGGCCGGGTCAGGTGTGGGAGGGGACACGTG | ATYGAGGTGAAGGYGAAGATAGYGAAGATTGTAGTGA GTGGGYGGGTYGGGTTAGGTGTGGGAGGGGATAYGTG | 7.5 |
| GCK | GGAGGGGGACGAGGGCCGCATGTGCGTCAATACCGAGT GGGG | GGAGGGGGAYGAGGGTYGTATGTGYGTTAATATYGAG TGGGG | 7.5 |
| TM6SF2 rs585429 C/T | CCA/GAGATCAGGCCTGCCT | No BS conversion | 7.5 |

##### 1.2. PDAL Seq protocol

PDAL-Seq was performed according to the protocol described in [1].

###### 1.2.1. Biotinylated P5 adapter

The following adapters were ordered in HPLC quality:

| Primer-ID | 5´mod | Sequence 5´-3´ | 3´Mod |
|---|---|---|---|
| P5-bio-top | Btn | ACACTCTTTCCCTACACGACGCTCTTCCGATCT | |
| P5-bottom | Phos | AGATCGGAAGAGCGTCGTGTAGGGAAAGAGTGT | 3InvdT |

###### 1.2.2. Biotinylated P7 adapter

The following adapters were ordered in HPLC quality:

| Primer-ID | 5´mod | Sequence 5´-3´ | 3´Mod |
|---|---|---|---|
| P7-top | Phos | GATCGGAAGAGCACACGTCTGAACTCCAGTCAC | dT-5' |
| P7-bottom | | GTGACTGGAGTTCAGACGTGTGCTCTTCCGATC | |

###### 1.2.3. Indexing PCR primers

*PCR primers:*

| Primer-ID | Sequence 5´-3´ |
|---|---|
| P5 primer (Universal) | AATGATACGGCGACCACCGAGATCTACACTCTTTCCCTACACGACGCTCTTCCGATCT |
| P7 primer (indexed) | CAAGCAGAAGACGGCATACGAGAT**NNNNNN**GTGACTGGAGTTCAGACGTGTG |

#### 1.3. Native DNA Polyacrylamide Gel Electrophoresis

##### 1.3.1. PCR primers

| Gene | Primer | Sequence | Length [bp] | Tm [°C] | Published |
|---|---|---|---|---|---|
| *MYC* Positive Control | Forward | TGGGCGGAGATTAGCGAGAG | 149 | 60 | [2] |
| | Reverse | CCTAGAGCTAGAGTGCTCGGC | | | |
| *ESR1* upstream enhancer region Negative Control | Forward | GAAACAGCCCCAAATCTCAA | 99 | 63 | [3] |
| | Reverse | TTGTAGCCAGCAAGCAAATG | | | |
| *GCK exon 7* | Forward | ATCCTTACAGCTGCTGACC | 227 | 60 | No |
| | Reverse | GCTCTCGCTGACAGTCC | | | |
| *TM6SF2 intron 2-exon 3* | Forward | TACCTCCTTGGTGTAGAACTCC | 197 | 54 | No |
| | Reverse | CCCTGACCTTAACTTCAAGCC | | | |

##### 1.3.2. Sample cleanup and reconstitution

PCR samples were cleaned up with ReliaPrep DNA concentration kit (Promega, Austria) and eluted in $dH_2O$. Concentrations were determined with Qubit HS DNA kit. Samples are reconstituted in a total volume of 20 µL with 50 mM Tris-HCl pH 8.3, or with 50 mM Tris-HCl pH 8.3 and 150 mM KCl with or without 10 µM pyridostatin (PDS, Merck, Austria) with a final concentration of 20 ng/µL.

##### 1.3.3. G4 folding protocol

G4 folding was carried out overnight in thermocycler according to the protocol below:

| | |
|---|---|
| 95°C | 7 min |
| 85°C | 10 min |
| 75°C | 60 min |
| 60°C | 60 min |
| 50°C | 60 min |
| 40°C | 60 min |
| 25°C | 60 min |

Ramp time 0.1°C/s

##### 1.4. Circular Dichroism (CD) Spectroscopy

The table below shows the oligos used for the CD spectra analysis. For *MYC* and *GCK* we have used the complement (G-rich strand), since the G4s are found in the -strands.

| Gene | Sequence |
|---|---|
| MYC | GGCCGACTCAGAGGAGGGGTGGAAGGGGTGGGAGGGGTGGGAGGGGTATTCGCGGGGAGGGCCCAAGGG |
| MYC-mutated | GGCCGACTCAGATGAGGTGTGGAAGGTGTGTGAGGTGTGTGAGGTGTATTCGCGGTGAGTGCCCAAGGT |
| MYC-methylated | GGC[5MedC]CGACTCAGAGGAGGGGTGGAAGGGGTGGGAGGGGTGGGAGGGGTATT[5MedC]CG[5MedC]CGGGGAGGGCCCAAGGG |
| GCK | GGGGCTTCCGCGGGGTGAGCCATAACTGCGTGTACGCCGGGAGCAGGGG |
| GCK-mutated | GGTGCTTCCGCGGTGTGAGCCATAACTGCGTGTACGCCGTGAGCAGGTG |
| GCK-methylated | GGGGCTTC[5MedC]CG[5MedC]CGGGGTGAGCCATAACTG[5MedC]CGTGTA[5MedC]CGC[5MedC]CGGGAGCAGGGG |
| TM6SF2 | GAGTGGGCGGGCCGGGTCAGGTGTGGGAGGGGACACGTGTGGGTGGGGCTAGGGCTTAGTGTGGGCGGGGCTTGAAG |
| TM6SF2-mutated | GAGTGAGCGGGCCGAGTCAGGTGTGAGAGAGGACACGTGTGAGTGAGGCTAGAGCTTAGTGTGAGCGAGGCTTGAAG |
| TM6SF2-methylated | GAGTGGG[5MedC]CGGGC[5MedC]CGGGTCAGGTGTGGGAGGGGACA[5MedC]CGTGTGGGTGGGGCTAGGGCTTAGTGTGGG[5MedC]CGGGGCTTGAAG |

##### 1.5. CpGfree Luciferase Reporter Assay Primers

| Gene | Primer | Sequence | RE site | Construct |
|---|---|---|---|---|
| *GCK* | Forward Cloning | tcagATGCATCGAACGGATTGTCAGTTTGC | NsiI | Promoter/Enhancer function (pCpGfree basic/promoter plasmid) |
| *GCK* | Reverse Cloning | tactGGATCCTTTCCATTGTTCCAGACAAAGC | BamHI | Enhancer function (pCpGfree promoter plasmid) |
| *GCK* | Reverse Cloning | tactAAGCTTTTTCCATTGTTCCAGACAAAGC | HindIII | Promoter function (pCpGfree basic plasmid) |
| *TM6SF2* | Forward Cloning | tcagATGCATGGTGTAGAACTCCATGAAGCC | NsiI | Promoter/Enhancer function (pCpGfree basic/promoter plasmid) |
| *TM6SF2* | Reverse Cloning | tactGGATCCGAATGAGGCAGGACTGTGG | BamHI | Enhancer function (pCpGfree promoter plasmid) |
| *TM6SF2* | Reverse Cloning | tactAAGCTTGAATGAGGCAGGACTGTGG | HindIII | Promoter function (pCpGfree basic plasmid) |
| Forward sequencing primer | | GCAGATTAAAAGGAATTCCTGC | | |

#### 2. Supplementary Tables

**Supplement Table 1. *P*-values of Study characteristics.** *P*-values are calculated with Students t-test

| | **LH vs LM** t.test *p* | **LH vs OH** t.test *p* | **LH vs OM** t.test *p* | **LH vs aMetD** t.test *p* | **LM vs OH** t.test *p* | **LM vs OM** t.test *p* | **LM vs aMetD** t.test *p* | **OH vs ON** t.test *p* | **OH vs aMetD** t.test *p* | **ON vs aMetD** t.test *p* |
|---|---|---|---|---|---|---|---|---|---|---|
| Age [y] | .3721 | .5957 | **.01524** | **1.53e-05** | .6934 | **.009013** | **9.98E-06** | **.01463** | **1.03E-05** | **.0006344** |
| BMI [kg/m2] | **.002851** | **< 2.2e-16** | **< 2.2e-16** | **4.56E-12** | **< 2.2e-16** | **< 2.2e-16** | **8.05E-10** | .7885 | **.007706** | **.005218** |
| HbA1c [%] | .3618 | **.004624** | **1.64e-05** | **2.93E-09** | .05583 | **0.0001127** | **4.93E-09** | **.003586** | **1.61E-08** | **3.53E-07** |
| HOMA | .1978 | **3.79E-05** | **2.33E-05** | **.01333** | **.000999** | **5.85e-05** | **.01507** | **.001521** | **.02342** | .07395 |
| FIB4 | .2752 | **.01954** | .3639 | **.0001389** | **.03671** | .8588 | **5.08E-05** | **.04789** | **1.84E-05** | **5.75E-05** |
| Fast. Glucose [mg/dL] | .3656 | **.04491** | **1.14E-05** | **2.54E-08** | .21 | **4.77E-05** | **3.78E-08** | **.0004376** | **6.08e-08** | **8.65E-07** |
| Fast. Insulin [mU/L] | .2375 | **4.40E-05** | **2.33E-06** | .05221 | **.00106** | **8.12E-06** | .06009 | .0007742 | .09915 | .261 |
| GGT [U/L] | **.006128** | **.01539** | **.004421** | **9.25E-05** | .1957 | .05061 | **.0001851** | **.01579** | **.000132** | **.0009559** |
| AST [U/L] | .1898 | .828 | **.01182** | **.0002775** | .2248 | .07171 | **.0005017** | **.0168** | **.0003004** | **.002234** |
| ALT [U/L] | .1018 | **.02687** | **.0001365** | **1.65E-06** | .5712 | **.001535** | **4.30E-06** | **.003457** | **6.20E-06** | **.0005368** |
| HDL-C [mg/dL] | .1161 | .2052 | **9.61E-05** | **1.18E-05** | .8108 | **.01587** | **.002933** | **.01113** | **.002114** | .5123 |
| LDL-C [mg/dL] | .7035 | .5417 | .4426 | **.02186** | .2872 | .623 | **.007109** | .1839 | .06205 | **.005796** |
| TG [mg/dL] | **.005892** | .1672 | **.0006377** | **3.69E-06** | **.03564** | **.04427** | **.00127** | **.001812** | **1.26e-05** | .4476 |
| TChol [mg/dL] | .646 | .5699 | .6297 | .1496 | .2283 | .9192 | **.04918** | .275 | .2847 | .06486 |

BMI=Body mass index, HOMA=homeostasis model assessment, $HbA_{1c}$=Glycated haemoglobin $A_{1c}$, FIB4=Fibrosis4 score, Fast. Glucose=fasting glucose, Fast. Insulin= Fasting insulin, GGT= γ-glutamyl transpeptidase, AST = aspartate aminotransferase, ALT = alanine aminotransferase, HDL-c= High density lipoprotein cholesterol, LDL-c= Low density lipoprotein cholesterol, TG= Triglycerides, TChol= total cholesterol

#### 3. Supplementary Figures

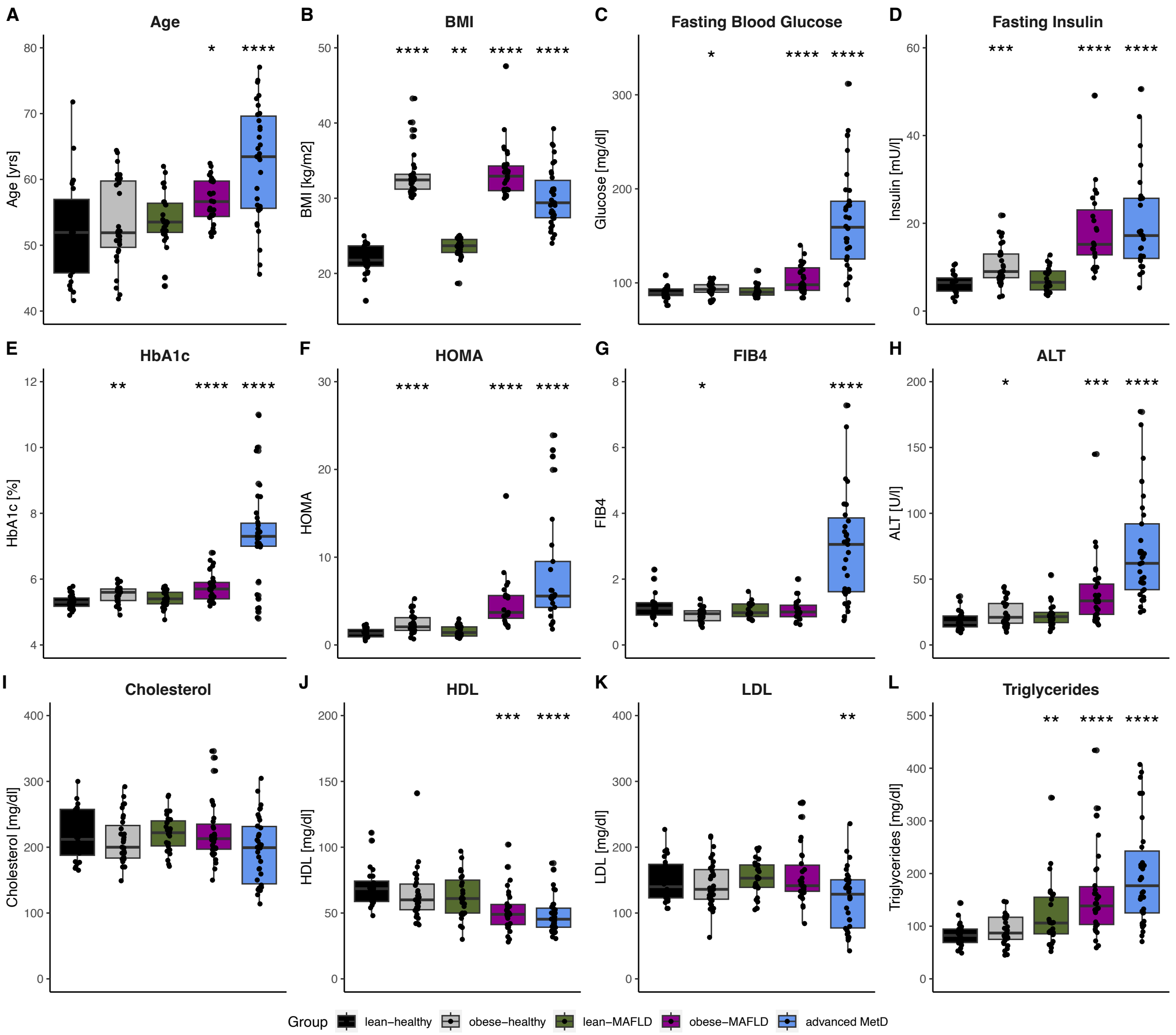

**Supplement Figure 1. Clinical Parameters of the study cohort.** The Box-and-whisker plots show the median of each group (black horizontal line) for all clinical parameters given in Table 1 of the main manuscript. Panel **A.** depicts the chronological age of the groups, **B.** the body mass index (BMI), **C.** Fasting Blood Glucose **D.** Fasting Insulin, **E.** glycated hemoglobin ($HbA_{1c}$), **F.** homeostasis model assessment (HOMA), **G.** Fibrosis 4 score (FIB4), **H.** alanine aminotransferase (ALT), **I.** Total Cholesterol, **J.** high density lipoprotein (HDL), **K.** low density lipoprotein (LDL), **L.** Total Triglycerides; Pairwise comparison against LH (t-test), *$p$<.05, **$p$<.01, ***$p$<.001, ****$p$<.0001, Exact $p$-values are listed in Supplement Table 1.

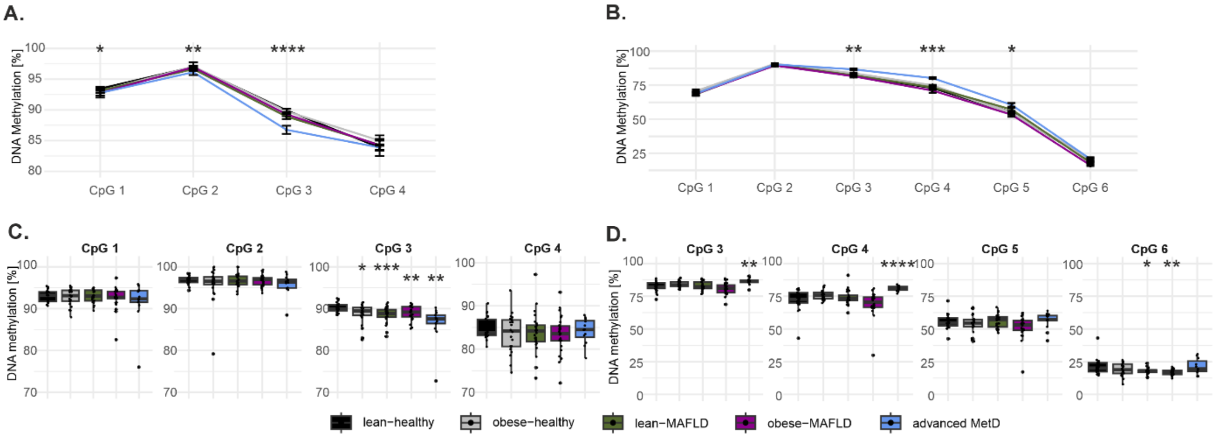

**Supplement Figure 2. DNA methylation in *GCK* exon 7 and the *TM6SF2* intron 2-exon 3 boundary.** Line plot showing the median DNA methylation in **A.** *GCK* over CpG 1-4 and **B.** in *TM6SF2* over CpGs 1-6. Stars (*) indicate the statistical difference between the advanced MetD and lean-healthy groups. DNA methylation at single CpG resolution by metabolic status for **C.** *GCK* exon 7 and **D.** *TM6SF2* intron 2-exon 3 for patients aged 42-60 years. Pairwise comparison against LH (Students t-test), *$p$<.05, ** $p$<.01, *** $p$<.001, **** $p$<.0001.

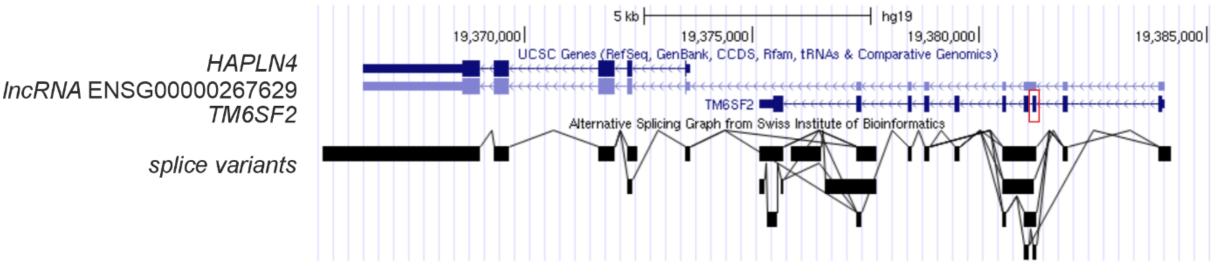

**Supplement Figure 3. *TM6SF2* alternative splicing variants extracted from the UCSC genome browser[4].** lncRNA ENSG00000267629 is formed by a readthrough from *TM6SF2* to *HAPLN4*. The cassette exon is marked with a red rectangle.

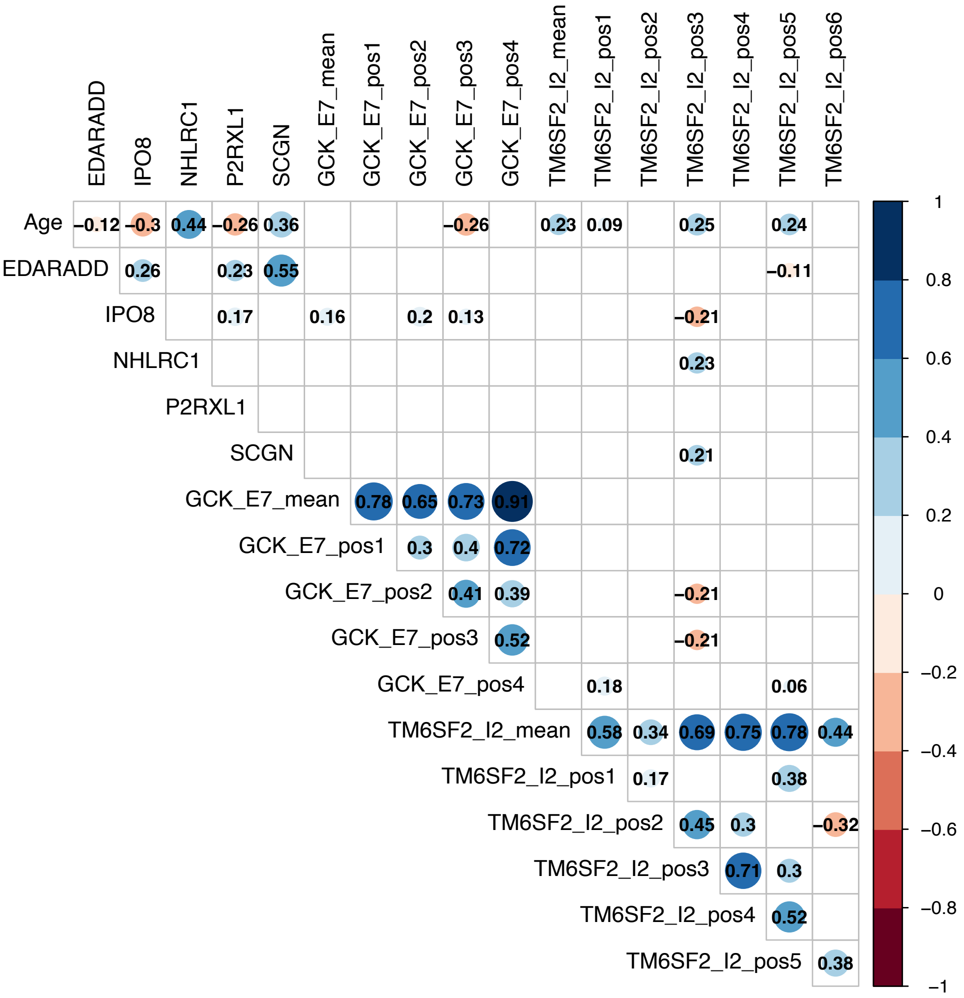

**Supplement Figure 4. Correlation Matrix investigating age and DNA methylation markers.** Published Epigenetic clock markers *EDARADD*, *IPO8*, *NHLRC1*, *P2RXL1* and *SCGN* detected by Illumina arrays [5-7] were investigated for DNA-methylation using locus-specific pyrosequencing assays. Their significant correlation with age was confirmed in our study cohort. Individual CpG positions of metabolism specific genes *GCK* exon 7 (CpG 3) and *TM6SF2* intron 2 (CpGs 1,3 and 5) as well as the mean DNA methylation over CpGs 1-6 in *TM6SF2* correlate with the blood donor's age. Only significant dependencies are shown. The color corresponds to positive (blue) or negative (red) correlations. Numbers in the circles represent the correlation coefficient and the size reflects p-values, with small circles less significant ($p$<.05) and big circles highly significant ($p$<.001). Nonsignificant relationships are blank.

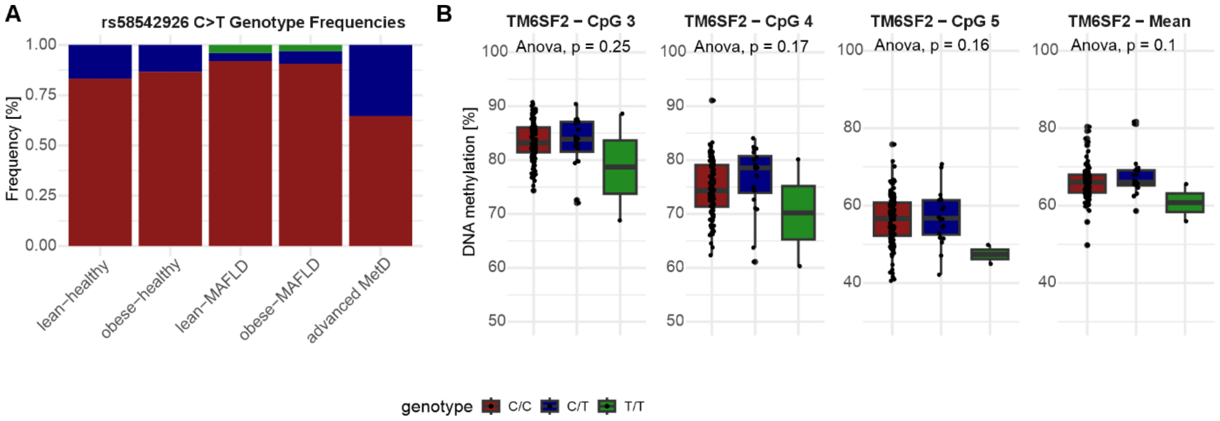

**Supplement Figure 5. Genotyping data for rs58542926 C>T. A.** shows the genotype frequencies in each group. **B.** DNA methylation values per genotype for CpGs 3-5, as well as the for the mean DNA methylation over the complete region (CpG 1-6); Statistics was calculated with ANOVA.

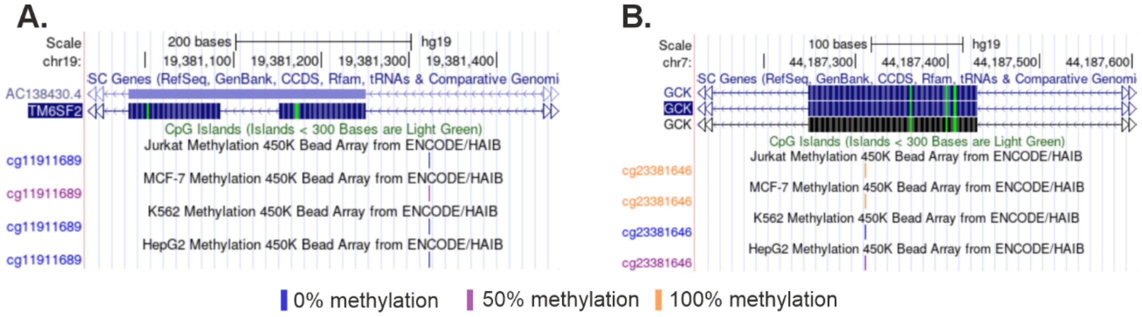

**Supplement Figure 6. DNA methylation of different cell lines at annotated CpGs from the Illumina Methylation array obtained from UCSC genome browser[4] in A.** *TM6SF2* and **B.** *GCK.*

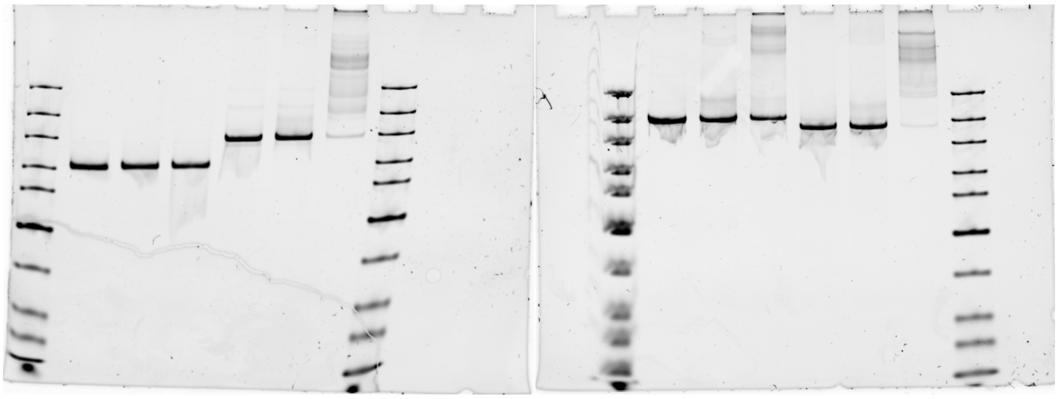

**Supplement Figure 7. Uncropped native PAA DNA gel of main figure 2B.**

#### 4. Supplementary Information

Within the TAD where *GCK* is located, several genes are known for their association with metabolic diseases and cancer. *AEBP1* was upregulated in MASH[8] and its elevated expression predicted poor prognosis of liver fibrosis[9]. *YKT6* was identified as metabolic syndrome risk gene in a genome-wide association study (GWAS) in the Taiwanese population[10] and the upregulation of *YKT6* transcription was associated with poor prognosis of HCC[11]. *BLVRA* expression was upregulated in HCC[12]. *URGCP* (also known as *URG4*) overexpression was related to tumor formation in mice[13], as well as in humans where it was found in several different cancer entities such as HCC, gastric and bladder cancer[14]. Overexpression of *PGAM2* was found to downregulate glycolysis genes like phosphofructokinase or hexokinase 2 in heart tissue of mice[15]. Two genes of the adjacent TAD were also linked to MetDs. The key cholesterol transport protein *NPC1L1* was identified as a risk factor for MAFLD[16]. *CAMK2B* differential methylation was caused by high glucose treatment in Huh-7 cells, which were used for MAFLD and T2D models[17]. Furthermore, interactive HiC analysis shows that *GCK* exon 7 contacts proximal enhancer region between *MYL7* and *POLD2.* Strikingly in a pan-cancer analysis, elevated POLD2 expression was found in advanced tumor stages and associated with poor prognosis, also for HCC[18].

*TM6SF2* is located in a TAD together with, a gene involved in targeting a lncRNA *UCF1* implicated in cancer formation and silencing of *UPF1* stimulated glycolysis in HCC[19]. Furthermore, a genetic variant located in the gene *NCAN* found in the same TAD is associated with MAFLD[20]. The interaction analysis showed that the *TM6SF2* DMP contacts a promoter-like region within intron 3 of *HPLN4* (~chr19:19,260,457, hg38) as well as intron 8 of *SUGP1* (~chr19:19,280,457, hg38), which was identified as regulator of cholesterol metabolism[21] (Fig. 3D).
